# Change point based dynamic functional connectivity estimation outperforms sliding window and static estimation for classification of early mild cognitive impairment in resting-state fMRI

**DOI:** 10.1101/2025.05.16.654552

**Authors:** Martin Ondrus, Ivor Cribben

## Abstract

The most widely used inputs in classification models of brain disorders such as early mild cognitive impairment (eMCI) or Alzheimer’s disease are estimates of static-based functional connectivity (SFC) and sliding window dynamic functional connectivity (swDFC). Although these methods are convenient for estimation and computational purposes, as it keeps the estimation tractable, they present a simplified version of a highly integrated and dynamic phenomenon. Change point dynamic functional connectivity (cpDFC) methods, which are far less commonly used, offer an alternative to swDFC approaches. In this study, we consider a classification task between controls and patients with eMCI using resting-state functional magnetic resonance imaging (fMRI) data from the Alzheimer’s Disease Neuroimaging Initiative (ADNI) studies, ADNI2 and ADNIGO. Our results indicate that the DFC methods are generally superior to the SFC methods when used as inputs into the classification model. Most importantly, we find that the cpDFC methods are generally superior to the widely used swDFC methods. We discuss how cpDFC methods offer many distinct advantages over swDFC methods, namely, the parsimony of network features and ease of interpretability. We validate the robustness and consistency of our results by testing the methods on an additional resting-state fMRI dataset of mild cognitive impairment patients. These findings call into question the validity of numerous fMRI studies that have utilized inferior approaches, such as SFC and swDFC, as inputs to classification models to predict various brain disorders. Finally, we present an ensemble model of the best models, which achieves an accuracy of 91.17% from leave-one-out cross-validation of subjects with eMCI. Our results suggest that the underlying functional networks are dynamic, multiscale, and that different FC methods capture distinct information for classification efficacy.

## 1 Introduction

Resting-state functional magnetic resonance imaging (rs-fMRI) is a non-invasive imaging technique with a simple protocol: patients are instructed to lie relaxed in the scanner for the duration of the session. Unlike task-based fMRI (tb-fMRI) studies, rs-fMRI does not require subjects to perform tasks, thereby reducing inconsistencies introduced by variations in experimental setup. Another key advantage of rs-fMRI is its ability to investigate interactions between brain regions over extended periods in a controlled and consistent environment. These interactions can reveal differences in connectivity patterns between healthy controls and individuals at the onset of brain disorders.

The primary goal of estimating functional connectivity (FC) is to quantify the statistical dependencies (Biswal et al., 1995) between regions of interest (ROIs) during a rs-fMRI or tb-fMRI experiment using measures such as coherence and correlation, among others (Cribben and Fiecas, 2016 provide a review). Graphs provide a natural framework for representing FC, where ROIs are depicted as nodes and their pairwise dependencies as edges. Graph-theoretic measures have been widely employed in neuroscience to describe the structural and functional organization of the brain (Muldoon and Bassett, 2016; Bassett and Bullmore, 2006, 2017), capturing the topological properties of the underlying network. FC can be further delineated into static (SFC) or dynamic (DFC), where time-dependent relationships are assumed to be either constant or changing, respectively. SFC has been used as an input in many state and clinical classification studies (Wee et al., 2012; Brier et al., 2012; Dai et al., 2012; Greicius et al., 2004; Sorg et al., 2007; Strain et al., 2022; Kim et al., 2024; Tu et al., 2024; van Nifterick et al., 2024).

Although SFC is simple to compute, it fails to measure the time-evolving relationships in the brain that have been shown to exist in both task-based (Fox et al., 2005; Eichele et al., 2008) and resting-state (Doucet et al., 2012) experiments. A diverse array of methods exist to estimate DFC, each offering unique advantages and limitations. The core premise of DFC is its ability to reveal how patterns of brain interactions evolve over time, providing insights into the temporal dynamics of neural activity. The sliding window approach is by far the most commonly used method for estimating DFC (Allen et al., 2014; Damaraju et al., 2014; Hutchison et al., 2013b; Hindriks et al., 2016) and has recieved a great deal of attention and extensive use in the literature. In this method, a time segment of a specified length is defined and progressively shifted forward across the time domain. At each shift, a new window is created and standard dependence measures, such as the correlation between regions of interest (ROIs), are applied to estimate FC.

The classification of psychiatric disorders using DFC measures has recently garnered interest, as DFC captures changing brain relationships from cognitive and state-related changes (see Du et al., 2018 and Filippi et al., 2019 for comprehensive reviews). Sliding window based DFC (swDFC) has been widely used in the literature to classify subjects by state or clinical conditions (Zhang et al., 2017; Schumacher et al., 2019; Zalesky et al., 2014; Karahanoğlu and Van De Ville, 2015; Chen et al., 2021; Kim et al., 2017; Canal-Garcia et al., 2024; Jing et al., 2023). However, the sliding window approach to estimating DFC has major limitations. Zalesky and Breakspear (2015) suggest fMRI patterns are stable in approximately 40 second intervals or longer, suggesting that changes in FC occur on the order of 10s of seconds. In the sliding-window method, the choice of window size is crucial: a window that is too large results in poor temporal resolution, while one that is too small yields less robust estimates (Cribben et al., 2012; Hutchison et al., 2013a). However, determining the optimal window size is often challenging and is typically unknown *a priori*. These decisions significantly influence the effectiveness of downstream tasks, as the windowing process determines the features used in the classification model, and is particularly critical in disease prediction.

The repetitive and overlapping nature of sliding windows has an additional drawback: it captures repeated and redundant connectivity patterns. In a high-dimensional multivariate time series setting such as fMRI data, this may lead to bias or erroneous variable selection in the classification model due to the large number of repeated windows and hence features. Individual snapshots that have been captured in sliding windows can be grouped together into “states” (or functional modes). However, the switching patterns between states, the number of states and the states themselves, may vary across individuals. This further confounds the implementation of sliding windows in a classification task. It remains unclear how to choose the number of states, whether they are shared within groups, or whether they generalize across all subjects regardless of disease status. It is plausible that each individual has a unique, time-varying “fingerprint” that may be similar to other subjects and different from other subjects.

To this end, in this work, we consider change point dynamic functional connectivity (cpDFC), as an alternative to SFC and swDFC methods, for extracting stable DFC features for early Mild Cognitive Impairment (eMCI) classification. Mild Cognitive Impairment (MCI) is a collection of precursor stages of AD that are inherently “mild” in both symptoms and brain morphology changes making them difficult to diagnose and detect in a clinical setting (Hampel and Lista, 2016). Characterizing differences between healthy controls (CN) and subjects with eMCI is of key importance to understand both healthy brain function and the neurodegenerative process. Given the lack of clear structural differences in the brain to identify eMCI, there has been interest in using functional imaging, in particular rs-fMRI, to understand these conditions (Petrella et al., 2011; Chen et al., 2017; Zhang et al., 2017).

In this study, we address a classification task that distinguishes controls from subjects with eMCI using rs-fMRI data from the Alzheimer’s Disease Neuroimaging Initiative (ADNI) studies, ADNI2 and ADNIGO. In the classification task, we compare the inputs of SFC, and swDFC across different window and step sizes to cpDFC methods. Unlike sliding windows, change point detection methods find non-overlapping stationary segments in the fMRI time series data, and is a generative, data-driven approach to find stable patterns across time series data, which reduces the redundancy of features. Furthermore, cpDFC methods can capture individual differences in dynamics by finding unique change points for each subject, whereas swDFC measures assume the same rate of change between subjects, which has been suggested to be an incorrect assumption (Leonardi and Van De Ville, 2015; Hutchison et al., 2013a; Hindriks et al., 2016). cpDFC methods applied to each subject may be more useful in uncovering parsimonious characteristics (or features) of DFC for disease classification.

Our work is unique and makes several key contributions. First, although many studies have explored DFC as input in brain disorder classification, to the best of our knowledge, this study is the first to explore the use of change point detection as inputs into a classification task of a neurodegenerative disease. Second, to be the best of our knowledge, no research has studied different step sizes for swDFC in a classification task. Third, our results indicate that the DFC methods (cpDFC and swDFC) are generally superior to the SFC methods when used as inputs in the classification model. Fourth, and most importantly, we find that the cpDFC methods, when used as inputs into the classification task, are generally superior to the extensively used swDFC methods. This finding calls into question the validity of numerous fMRI studies that have utilized inferior approaches, such as SFC and swDFC, as inputs for classification tasks for various brain disorders. Fifth, by comparing classifier performance and the selected inputs across the SFC, swDFC and cpDFC approaches, we find that cpDFC provides strong consistency and interpretability in this complex and challenging setting. Sixth, we show how classification of eMCI using rs-fMRI data benefits greatly from understanding both local and global brain functional dynamics, which we characterize through the estimation of graph theoretic measures. Seventh, we validate the robustness and consistency of our results by testing the methods on an additional resting-state fMRI dataset of subjects with mild cognitive impairment. Eighth, we present an ensemble model of the best methods (SFC, swDFC and cpDFC), which achieves an accuracy of 91.17% from leave-one-out cross-validation of subjects with eMCI. Typical classification results range from 70% to 90% depending on the modality. Our results suggest that the underlying functional networks are dynamic, multiscale and that different FC methods capture distinct information for classification efficacy. In general, our work underscores the versatility and practicality of change point detection in fMRI experiments, particularly in its application to downstream tasks such as classification. We also hope that this study inspires renewed interest in advancing research on change point detection techniques aimed at estimating dynamic functional connectivity.

The remainder of the paper is organized as follows. In Section 2, we detail the data and preprocessing steps. In Section 3, we describe the notation, the static based, the sliding window and the change point detection methods. We also define the graph topological features that are used to summarize the networks and the classification models. We outline our classification study in Section 4 and present our results in Section 5. In Section 6, we discuss our findings and conclude in Section 7.

## 2 Data and pre-processing

### 2.1 ADNI data

The first dataset we obtained from the ADNI database (http://adni.loni.usc.edu). ADNI was launched in 2003 as a public-private partnership, led by Principal Investigator Michael W. Weiner, MD. The primary objective of ADNI has been to test whether serial magnetic resonance imaging, positron emission tomography, other biological markers, and clinical and neuropsychological evaluations can be combined to measure the progression of MCI and early AD. For up-to-date information, see www.adni-info.org. In ADNI’s resting-state functional magnetic resonance imaging (rs-fMRI) experiments, subjects were instructed to remain still and relaxed in the scanner. Subjects level data were obtained using Phillips scanners and include 33 subjects with eMCI (mean age 72.3, 15M/18F) and 35 healthy controls (mean age 74.6, 14M/21F).

Data were pre-processed using SPM (SPM, 2023), following a standard fMRI preprocessing pipeline. The first 10 volumes were discarded to account for initial scanner and subject noise. Next, a slice timing correction was performed (`spm_slice_timing`) to correct for timing differences between slices within each volume. The images were then realigned using `spm_realign`, which applies a rigid-body transformation to align each volume with the mean functional image. Motion parameters were estimated using least-squares with 2nd degree B-spline interpolation. The estimated motion parameters were subsequently applied to reslice all volumes using 4th degree B-spline interpolation to minimize resampling artifacts. The images were then normalized to the Montreal Neurological Institute (MNI) space with 3mm×3mm×3mm voxels (`spm_normalise`). Nuisance covariates (Friston 24, cerebrospinal fluid, white matter, and global mean) were regressed out. The voxels were then spatially smoothed using a Gaussian kernel (FWHM = 6mm) in `spm_smooth`. A linear trend was removed from each time series using ordinary least squares regression (`spm_detrend`) and a fourth order Butterworth low-pass filter (0.01–0.08 Hz) was applied to attenuate high-frequency noise and low-frequency drift outside the typical BOLD signal range (`spm_filter`). We used the Automated Anatomical Labeling (AAL2) atlas to subdivide the brain into 120 anatomical regions (Tzourio-Mazoyer et al., 2002). The region of interest (ROI) time series for a given region was defined as the average of the time series of all voxels within that region. Finally, each time series was z-score normalized. The final preprocessed dataset for each subject was a time series of dimension *T* = 130 time points by *p* = 120 ROIs.

### 2.2 Second MCI data set

In order to check the robustness of our results, we apply our methods to a second study of MCI classification (Mascali et al., 2015), which includes 10 patients with mild cognitive impairment and 10 healthy elderly controls. Participants were told to lie quietly with their eyes closed without falling asleep. A 3T MRI system (Magnetom Allegra, Siemens, Erlangen, Germany) was used to acquire images, with the following properties: TR = 2080 ms, TE = 30 ms, 32 axial slices parallel to the AC-PC plane, matrix = 64 x 64, in plane resolution = 3x3 mm^2^, slice thickness = 2.5 mm, 50% skip, flip angle = 70^°^. Functional images were preprocessed using the Connectivity toolbox (Whitfield-Gabrieli and Nieto-Castanon, 2012). The initial four volumes were discarded for signal and scanner stabilization, resulting in 216 time points per subject, and images were slice-time corrected and realigned to the first image. More detailed information on the preprocessing steps can be found in Mascali et al. (2015). Finally, the atlas of Gordon et al. (2016) was used to parcellate into ROIs, and then the Default (Sperling et al., 2010; Buckner et al., 2008), Frontoparietal (Brier et al., 2012), Cinguloparietal (Bai et al., 2009), and Dorsal Attention (Zhou et al., 2008) communities were selected to reduce the dimensionality to 102 ROIs, in order to be comparable to the main study, which has 120 ROIs.

## 3 Methods

### 3.1 Notation

We provide the notation used for the remainder of the paper. We denote a matrix with a bold capital letter ***A*** and vectors as bold lowercase letters ***a***. The entry corresponding to the *i*th row and the *j*th column in ***A*** is ***A***_*ij*_. The vector of the *i*th column in ***A*** is given by ***A***_*i*_. We denote a single subject’s fMRI data by ***X***, where each data matrix ***X*** ∈ ℝ^*T* ×*p*^ has *T* time points and *p* ROIs. A time point *t* is an element of the time index set {1, …, *T*}. A stationary segment in ***X*** is denoted by ***S*** = {*x*_*t*_ ∈ ℝ^*p*^: *t*_1_ ≤ *t* ≤ *t*_2_} where *t*_1_, *t*_2_ ∈ {1, …, *T*}. An unweighted graph 𝒢 is defined as a collection of vertices and edges 𝒢 = (*V, E*). We use the terms “graph” and “network” interchangeably throughout.

### 3.2 Functional connectivity through graph estimation

From the DFC approaches outlined in the next section, we obtain a series of stationary segments for each subject, while for SFC, stationary is assumed over the entire experimental time course, and thus one segment is obtained for each subject. For each stationary segment ***S***, we estimate the FC or linear dependency between the *i*th and *j*th time series (or vertices) in 𝒢 using Pearson’s correlation coefficient, *ρ*_*ij*_. We can threshold 𝒢 by applying a cutoff. Although there is no universal agreement on the best approach, most studies typically apply a cutoff in the range of 0.1 to 0.8 (Adamovich et al., 2022). We seek to find stability; that is, we select a cutoff value which produces graphs that are not overly dense and difficult to interpret, or not overly sparse thereby losing important information about connectivity. We are interested in strong correlations; hence an edge *E* in 𝒢 between the vertices *i* and *j* remains in the graph if |*ρ*_*ij*_| *>* 0.5. A schematic of SFC estimation is shown in Figure 1(b).

**Figure 1:**
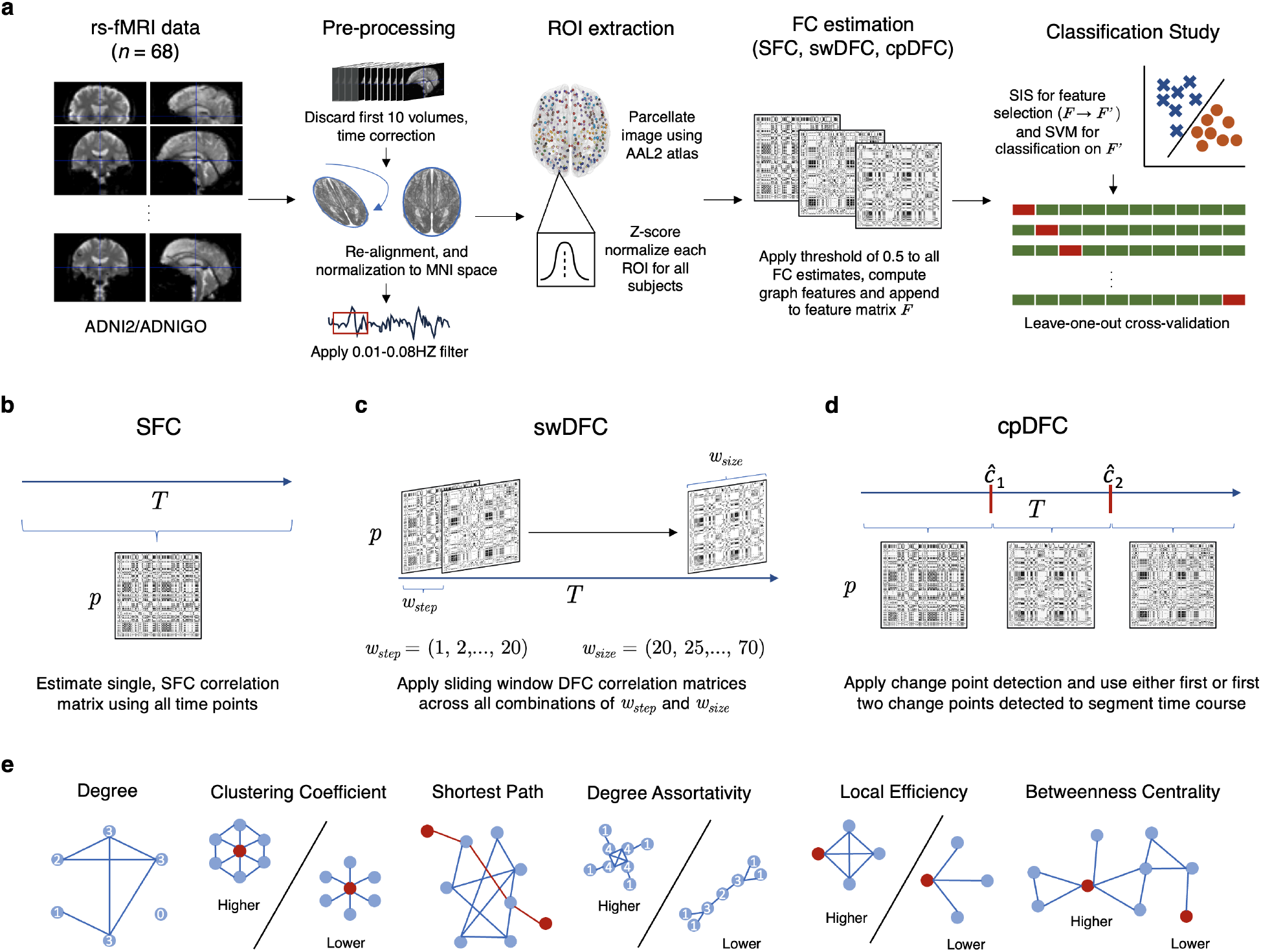
A schematic of CN/eMCI classification task starting from raw rs-fMRI data to eMCI/control classification for the ADNI rs-fMRI dataset. Panel (a) is the overall pipeline, while the middle three panels (b), (c), and (d) show the different FC methodologies (SFC, swDFC, cpDFC, respectively). The bottom panel, (e), shows example graphs expressing the topological features used in the classification study. Nodes are labeled with their degree, and the color red is used to highlight the nodes and edges of interest.

### 3.3 Estimating DFC

To estimate DFC, we assume that we have no prior knowledge of the distribution of ***X***, and that there may be an unknown number of distributional shifts, with the number and locations in {1, …, *T*} and the rate of change between them is assumed unknown. We do however assume that there exist segments in ***X*** which *are* stationary. We consider two approaches – sliding windows (swDFC) and change points (cpDFC) – for estimating time-varying connectivity in rs-fMRI. swDFC is a naive approach in which no data information is used to determine the temporal structure. Here, a window of a particular length, *w*, is used to subselect the set of time indices in {1, …, *T*}. As its name suggests, the window is then slid over a predetermined length, *s*, to define the next window. Consequently, the sliding window captures dynamic information by subsampling the entirety of the distribution through a predetermined and overlapping set of time-dependent steps, where within each of these windows, FC is estimated as described in Section 3.2. We describe the sliding-window method in Algorithm 1 (in the Appendix) and show a schematic in Figure 1(c).

The choices for *w* and *s* are crucial, as they determine the granularity and the amount of overlap between snapshots that this technique captures. However, the values for these parameters are context dependent and it is not possible to derive them directly from ***X***. As a consequence of this, we vary the combinations of *w* and *s* in our experiments and test the classification performance as described in Section 4.

We implement cpDFC using FaBiSearch, a change point detection technique in the network structure between (high-dimensional) multivariate time series (Ondrus et al., 2025). Change point detection is a commonly used method to find abrupt changes in the distribution of time series data. More specifically, we consider the multiple change point detection problem in a multivariate time series dataset ***X***. We seek to find the time points where the network structure of ***X*** changes. FaBiSearch utilizes non-negative matrix factorization (NMF: Lee and Seung, 1999). Algorithm 2 (in the Appendix) summarizes FaBiSearch, and we utilize a loss based on Kullback-Leibler divergence to assess the fit of the model. For more details on FaBiSearch, see Ondrus et al. (2025).

In contrast to the sliding window method, the inputs into FaBiSearch need only be *sufficient* rather than *optimal*. Ondrus and Cribben (2024) show through a sensitivity analysis on simulated data that the accuracy of change point detection using FaBiSearch plateaus beyond sufficient values of the input hyperparameters. In particular, they show that *rank* can be found effectively before the change point procedure, and FaBiSearch achieves optimal performance when *δ* ≥ 30, *n*_*run*_ ≥ 100, and *n*_*reps*_ ≥ 100, which denote the minimum distance between change points, the number of runs for NMF convergence, and the number of permutations in the hypothesis test, respectively.

For each subject, we first estimate the change points using FaBiSearch and then order the change points 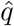 by their respective *p*-value from smallest to largest. After defining the set of change points, 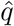, we partition ***X*** into stationary time segments between the change points. In the classification study, we used *k* = 1 and *k* = 2 change points to define the stationary segments. Finally, for each of the stationary segments, we estimate FC as described in Section 3.2.

We also estimate cpDFC using two other change point detection methods, specifically Network Change Point Detection (NCPD: Cribben and Yu, 2017) and Covariance Change Points through Random Matrix Theory (CRMT: Ryan and Killick, 2023). For NCPD and CRMT, we used similar hyperparameters to FaBiSearch, that is, a similar minimum distance between the change points and the optimal rank. All other hyperaparameters for these methods were set to their default values. For CRMT, we further pre-processed ROI time series by performing a truncated SVD with the same optimal rank as used in FaBiSearch to satisfy the theoretical condition that *p* ≲ *n* for the estimator to be well-behaved. A schematic of change point detection is shown in Figure 1(d).

### 3.4 Graph topological features

To classify each subject, we use the topological characteristics of the graphs of each stationary segment. For each graph 𝒢, we generate features that incorporate a wide variety of node-wise graph properties, and then use variable selection with an internal cross-validation procedure to select the best combination for classification, which is a common procedure (Gheiratmand et al., 2017). We now describe these features and include a visual description in Figure 1(e), with formal definitions provided in the Appendix.

Degree is a local graph summary metric and is calculated for each node by counting the number of edges that connect to it. Those with high degree are typically perceived as more important and have “hub-like” behavior (Wasserman and Faust, 1994), as they tend to integrate more information from other nodes. Decreases in the degree of important nodes may reflect losses in the integration of information from other brain regions, and thus defects in neural processing abilities (Sporns, 2016). The clustering coefficient is a local graph summary metric that quantifies the proportion of connected adjacent nodes. A higher value indicates more dense, interconnected groups of adjacent nodes (Wasserman and Faust, 1994). It is typically involved in the small world hypothesis of brain organization and can be useful in understanding node localization and integration (Watts and Strogatz, 1998). In the context of network neuroscience, differences in node clustering coefficient implies a functional organization where a few nodes integrate information from many others, and can highlight differences between the functional roles of brain ROIs (Rubinov and Sporns, 2010; Sporns, 2016). The shortest path is also a local graph summary metric denoting the smallest number of edges it takes to go from one node to another in the graph, calculated for each unique pair of nodes in the graph. It can be interpreted as the inverse of the relative ease of information transfer across the network, where a higher value indicates more difficulty and vice versa (Rubinov and Sporns, 2010). Increased shortest path metrics indicates poorer information transfer (Rubinov and Sporns, 2010). The degree assortativity is a global graph summary metric, which increases if nodes with a high connectivity are connected to other nodes that also have high connectivity (Newman, 2002). A large assortativity value describes a network with a “rich club” where the highly interconnected nodes are typically connected to each other (Colizza et al., 2006). This coefficient describes the extent to which important hub nodes are interconnected to each other, and can be a useful property when trying to investigate and understand the brain’s resilience to functional declines (Van Den Heuvel and Sporns, 2011). The local efficiency is a local graph summary metric, wherein each node is removed from the graph, and the average shortest path length between the nodes in the rest of the network is calculated. The local efficiency is then the inverse of this, and denotes how important each node is in interconnecting other nodes in the network (Latora and Marchiori, 2001). This measure shows how effective functional wiring of the brain is and how quickly and effectively information can be disseminated (Bassett and Bullmore, 2006; Achard and Bullmore, 2007; Bullmore and Sporns, 2009). The betweenness centrality is a local graph summary metric and is measured by the frequency of shortest paths that go through that node. Thus, it measures the centrality of that node for the traversal between different parts of the graph (Freeman, 1977). In the context of network neuroscience, Bassett et al. (2010) considered betweenness centrality through the lens of a circuit, and concluded that the wiring of the brain is heavily dependent on the routing of information, and the efficiency of transfer which is reflected through this measure. Additionally, Crossley et al. (2014) described betweenness centrality in the context of brain disorders, implying regions that are typically high in this measure may be inefficient or defective in those subjects with neurodegenerative disorders.

### 3.5 Feature selection and classification

We define class labels *c* = 0 for the control group and *c* = 1 for the eMCI group. Our objective is to compare and contrast SFC, swDFC, and cpDFC separately as inputs into a classification model. For the DFC methods, each subject has feature vectors from each stationary segment which are concatenated column-wise. Before variable selection, the feature vectors are concatenated for each subject into the feature matrix ***F***. Although computing graph features markedly reduces the number of input features in the classification model, compared to using the raw edge weights, there are still many more features than samples (*p >> n*). We use sure independence screening (SIS), which uses correlation learning to efficiently find predictive features and also has particular properties that make it well suited to a high dimension classification setting (Fan and Lv, 2008). The final feature matrix after variable selection is ***F*** ^′^.

We used a linear SVM model (Boser et al., 1992) on the final feature matrix ***F*** ^′^ to predict the *c* classes. In addition to linear SVM, we also tested RBF-SVM, logistic regression, and decision trees. In all studies, we found the linear SVM to be the most robust and thus we choose to focus on this model, as it provides strong theoretical and practical implications (Suykens and Vandewalle, 1999; Boser et al., 1992).

## 4 Classification Study

After pre-processing the data, we split the study into three tasks based on the type of FC measures used: SFC, swDFC, and cpDFC. For the SFC task, all the *T* = 130 experimental time points were used to calculate the FC. For the swDFC task, we used the sliding window technique described in Section 3.3 and tested a variety of different windows (time window of length 10 to 70 with increments of 5) and step sizes (1, 2, 3, 5, 8, 10, 15, 20). Lastly, for FaBiSearch, for each subject, we ordered the detected change points based on their *p*-value. The first change point (FBS cpDFC1) or the first two change points (FBS cpDFC2) were used in the final model. SIS for feature selection was implemented using Saldana and Feng (2018), followed by classification using a linear SVM from Meyer et al. (2024). Training and testing was carried out using leave-one-out cross-validation on the 68 subjects. Feature selection through SIS and training of the classifier model were repeated independently within each fold. These steps are summarized in Figure 1(a).

We also explored an ensemble method, which combines the best classifiers, in order to determine whether additional information can be gained by combining the SFC, swDFC, and cpDFC approaches. In particular, we combined the best 2 models, 3 models, …, 10 models (as determined by cross validation F1 score) by taking the average of prediction probabilities that were calculated within each fold and then testing using leave-one-out cross validation, for the individual FC condition models. We formally detail all evaluation metrics used to assess models in the Appendix.

## 5 Results

### 5.1 Change point detection and stationary segments

To estimate cpDFC, we apply FabiSearch to each subject in the ADNI dataset. Figure 2 shows both the detected change points for the control group (CN) (Figure 2(a)) and the eMCI group (Figure 2(b)). There does not appear to be a difference in the pattern or the order of the change points between the two groups, suggesting that the change points are unique across subjects. The detected change points are distributed quite uniformly for both groups, although there do appear to be some edge effects with some change points being concentrated near the beginning and end of the scanning session. A non-parametric Kolmogorov-Smirnov test on the change point locations between the two groups suggests that they are not statistically significantly different (*p* = 0.413). A one-sided *t*-test on the means of the location of the first detected change points of the CN group and the eMCI groups suggests that the first change point for the CN group is statistically significantly smaller (in time) than the eMCI group (*p* = 0.028).

**Figure 2:**
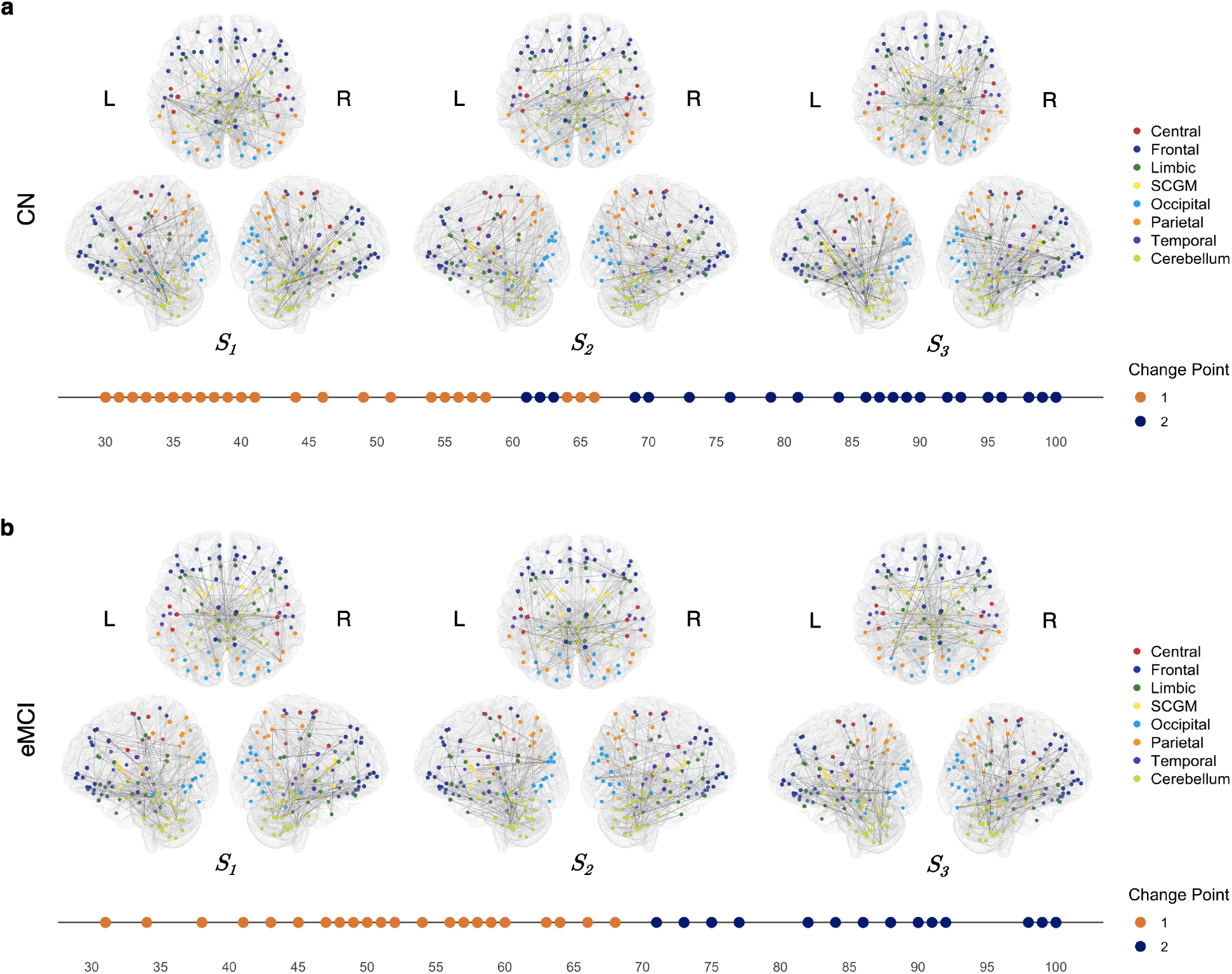
The change point detection results from applying FaBiSearch (taking only the first two change points, FBS cpDFC2) and corresponding stationary FC states for (a) controls and (b) subjects with eMCI for the ADNI rs-fMRI dataset. For each panel, FC plots are shown for each stationary segment, where ***S***_1_, ***S***_2_, ***S***_3_ correspond to the first, second, and third stationary segments. Individual change points are used to segment the time series, and then correlation matrices are averaged across subjects. The top 100 edges as determined by the absolute value of this averaged correlation are shown for each stationary segment. Below the FC plots, we depict the location of the first two change points detected across all subjects in each group.

Figure 2 also shows the stationary FC states (or modes) between each change point. Across all segments and even between the CN and subjects with eMCI, there is strong FC between the frontal and cerebellar regions. In the first FC state, ***S***_1_, CN have stronger FC between the parietal regions and the rest of the brain, and also engage the temporal regions with the rest of the brain more as well. Subjects with eMCI, have stronger FC between the frontal and parietal regions.

In ***S***_2_, CN subjects exhibit strong FC between the frontal and parietal regions. In contrast, subjects with eMCI show FC primarily concentrated in the frontal regions, with connections extending to the occipital and limbic systems. Notably, a small number of occipital nodes in subjects with eMCI mediate much of the FC to other brain regions. In ***S***_3_, CN patients display robust FC among the frontal, occipital, and cerebellar regions, forming a tightly coupled triad. Conversely, subjects with eMCI exhibit more diffuse FC, with the cerebellum acting as a hub, extensively connected to the rest of the brain – particularly the parietal lobe.

### 5.2 Classification

We first evaluate the SFC, swDFC, and cpDFC approaches separately, and then we evaluate an ensemble of classifiers with the best-performing models. Figure 3 shows the main results of our study. Due to the large number of window and step size combinations, we limit the swDFC results to only the combinations that achieve an accuracy of at least 63.57% (which is one standard error above the null accuracy of 51.47%). Of the SFC, swDFC and cpDFC methods, change point detection with two change points (cpDFC2) has the best overall performance in three of the four performance measures (see the Appendix for their definitions): accuracy 77.94%, F1 score 80.00%, sensitivity 90.91%, and specificity 65.71%. The only measure of performance where FBS cpDFC2 was inferior to swDFC was sensitivity. In this case, the two combinations of swDFC that outperformed FBS cpDFC2 (60w 15s and 70w 10s) in terms of sensitivity both performed poorly compared to FBS cpDFC2 on the other performance measures. Compared to the best swDFC combination, swDFC 15w 3s (accuracy; 72.06%, F1 score; 74.67%, sensitivity; 84.85%, specificity; 60.00%), FBS cpDFC2 only marginally outperformed it (*p* = 0.250). However, in the set of all combinations of swDFC (*µ* = 0.5852, *σ* = 0.0483), FBS cpDFC2 had statistically significant superior performance in the four main performance measures (accuracy; *p* = 2.12 × 10^−6^, F1 score; *p* = 1.38 × 10^−5^, sensitivity; *p* = 1.19 × 10^−6^, specificity; *p* = 6.18 × 10^−6^). cpDFC1 performed poorly compared to FBS cpDFC2 and the best performing swDFC methods. In general, the SFC performed poorly. In particular, it consistently performed close to the null performance of this dataset, 51.47%. While SFC has poor performance, it was at least stable, compared to the swDFC methods which had greater variability in performance.

**Figure 3:**
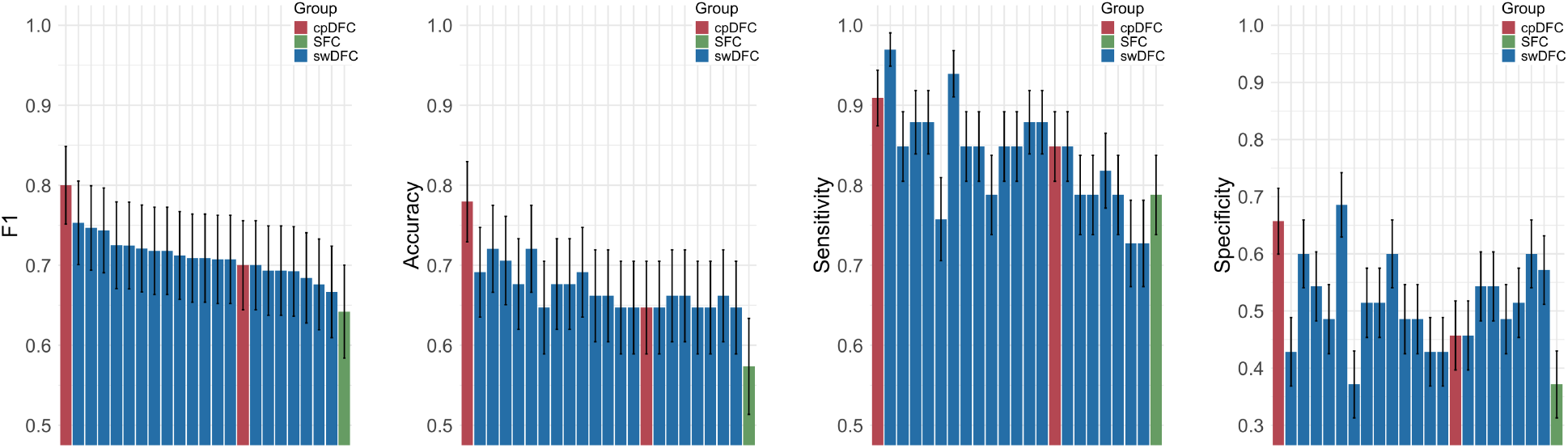
F1, accuracy, sensitivity, and specificity results from the classification study of CN subjects and subjects with eMCI in the ADNI rs-fMRI dataset. SFC, cpDFC, and swDFC correspond to static, change point, and window based dynamic functional connectivity, respectively.

We also consider the number of unique features that were selected across the 68 folds for all methods, where a lower number of unique features indicates greater stability as there is more consistent selection of the same features across folds. Here, FBS cpDFC2 chose less than the swDFC models, which is a consequence of less variability in the features chosen using the SIS method. We find cpDFC offers good stability, with 28 unique features across all folds, with SFC having less stability, although still adequate at 37. In contrast, the swDFC methods are not as consistent. There is no clear relationship between the stability of the selected features and the window size and the step size. Furthermore, although the best four swDFC methods outperform cpDFC1 and FBS cpDFC2, the differences are marginal. Furthermore, for swDFC, there does not seem to be a clear relationship between model performance and feature selection stability, which suggests a trade-off between model performance and stability for the swDFC approach. Figure A3 shows the unique features selected across LOOCV folds for FBS cpDFC, swDFC, and SFC.

Figure 4 shows the results of different combinations of window size and step size for swDFC. Across the various combinations, there is no clear relationship between step size and window size. Additionally, there are several combinations that are plagued by instability, as shown by the large variability in step and window sizes in neighboring combinations (or tiles). For example, the combination of window size 15 and step size 3 achieved the best accuracy of 72.06%. However, changing the window size by 5 time points in either direction (to either 10 or 20), drastically reduces the performance of these models (to 58.82% and 55.88%, respectively). The same is also evident when the step size is changed to 2 or 8: the performance of the models reduces to 52.94% and 64.71%, respectively.

**Figure 4:**
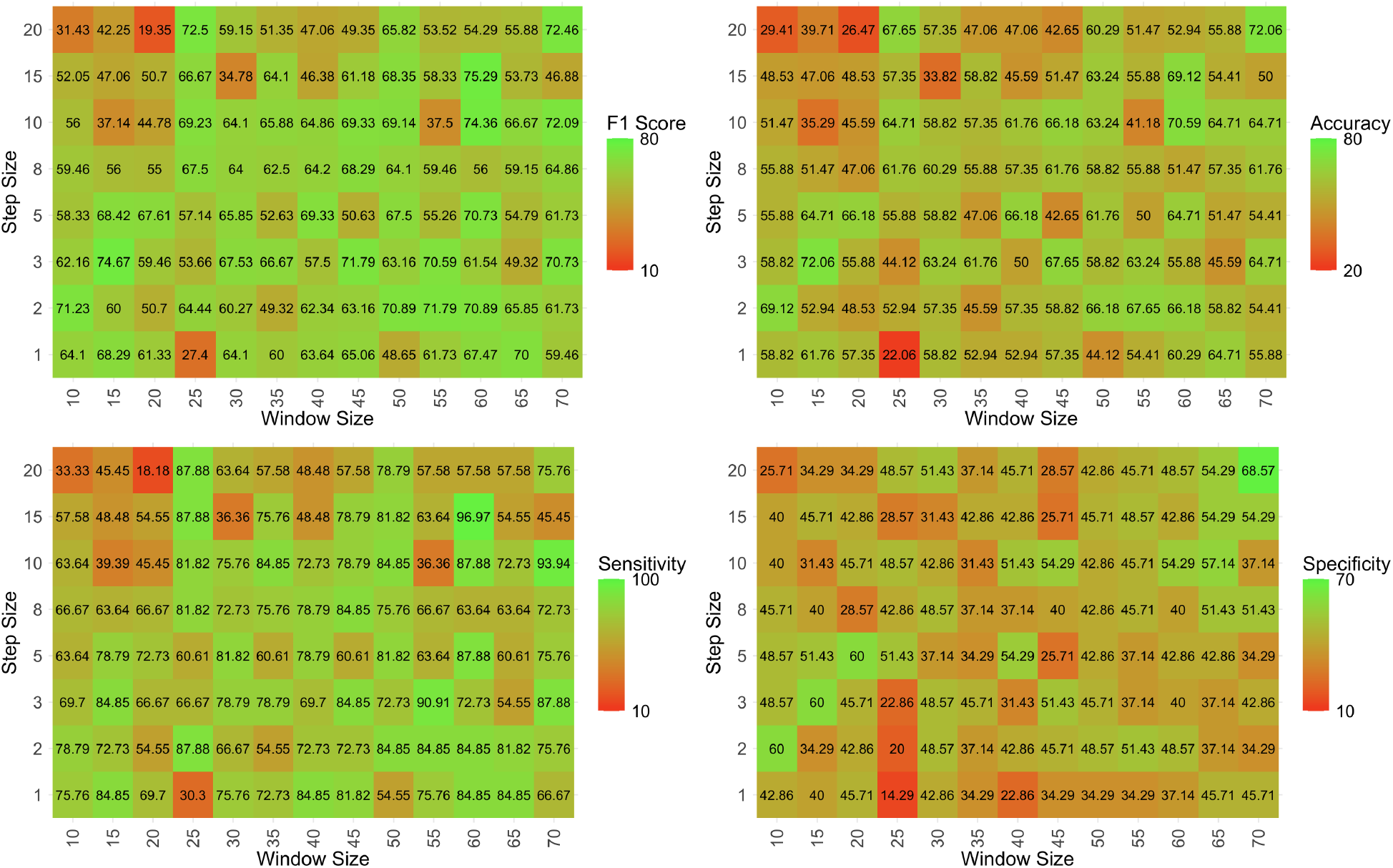
Heatmaps of F1 score, accuracy, sensitivity, and specificity from the classification study of CN subjects and subjects with eMCI from the ADNI rs-fMRI dataset using swDFC. Results are shown across window sizes [10, 70] in increments of 5 and step sizes [1, 2, 3, 5, 8, 10, 15, 20].

In Figure 5(a), we show the ROIs that were selected by SIS in all folds in leave-one-out cross-validation for FBS cpDFC2. Figure 5(b) includes more information on the selected features and the corresponding ROIs, such as node ID, feature type, stationary segment, and differences in the mean value for the features of the CN and the eMCI groups. We find that the most consistently chosen features correspond to the frontal, parietal, and cerebellum regions. Degree, betweenness, clustering coefficient, local efficiency, and shortest path were all key features that were selected across LOOCV folds. We also find that eMCI is associated with lower degree in the parietal region, and that paths were more efficient, shorter, and tightly clustered in the cerebellum and frontal regions. Later segments (***S***_2_, and ***S***_3_) were chosen more often across these folds, which may be related to the subjects being more settled and closer to a true “resting” state compared to the beginning of the fMRI experiment.

**Figure 5:**
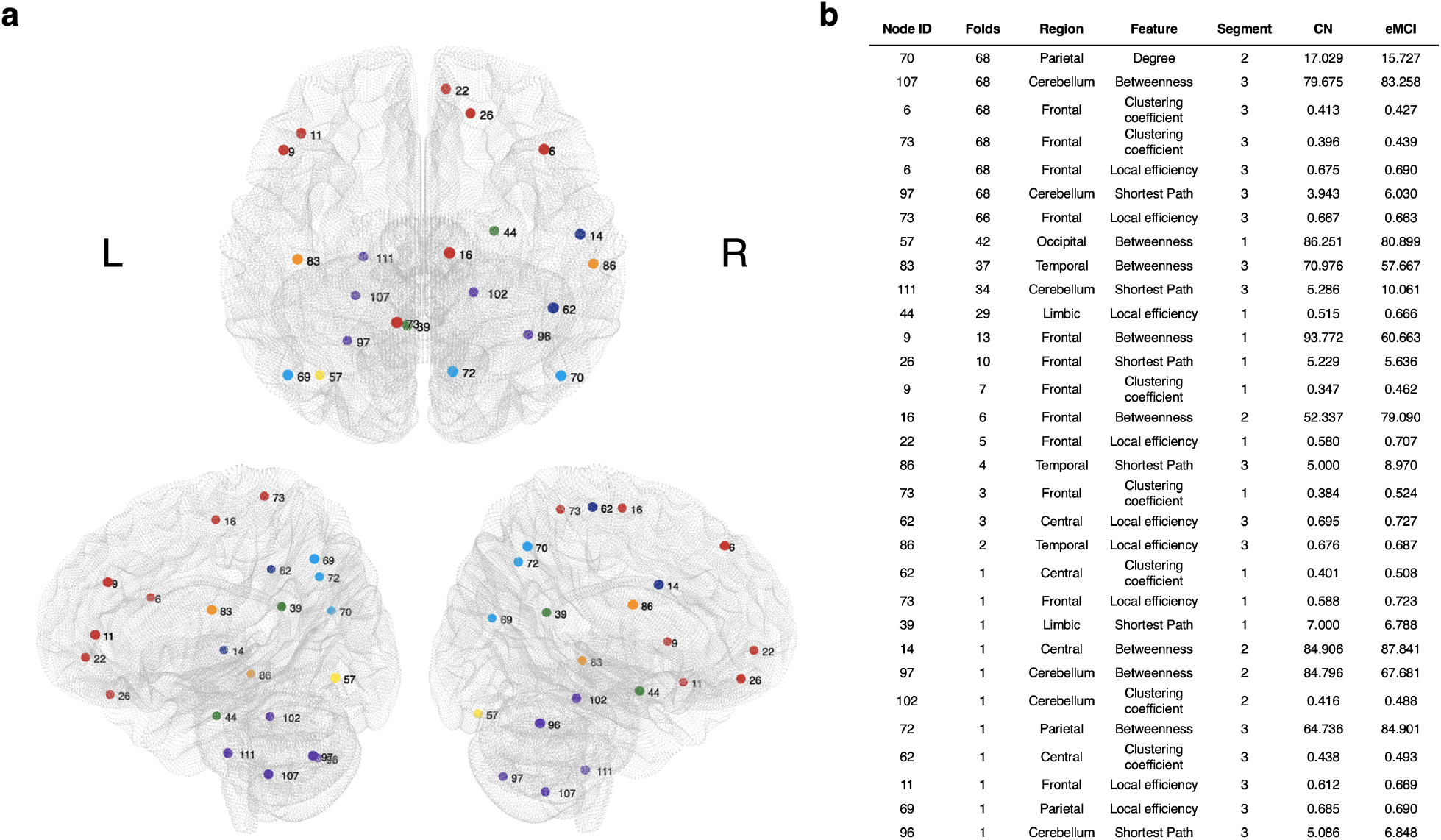
(a) The associated regions of interest (ROIs) of the features selected across all folds by SIS in leave-one-out cross validation for FBS cpDFC2 in the classification study of CN subjects and subjects with eMCI from the ADNI rs-fMRI dataset. (b) The node ID, the number of LOOCV folds that the feature was chosen, region from the AAL atlas, the graph theoretic feature type, the stationary segment (i.e., 1 = first, 2 = second, 3 = third), and the mean values of the features of the CN and eMCI groups. The features are ordered in descending order based on how often they were selected across the LOOCV folds.

### 5.3 Additional change point methodologies

To compare the performance of FaBiSearch, we also applied both the NCPD and the CRMT change point methods to the ADNI rs-fMRI dataset. Figure 6 shows the detected change points for each method while Table 1 show the classification results. We find that FaBisearch with two change points, again, has the best performance. CRMT with one change point has the next best performance in terms of accuracy, followed by NCPD. NCPD with one change point has the worst accuracy. Similar to FaBiSearch, we find that two change points perform better than one change point in all change point methods, in terms of accuracy.

**Table 1:**
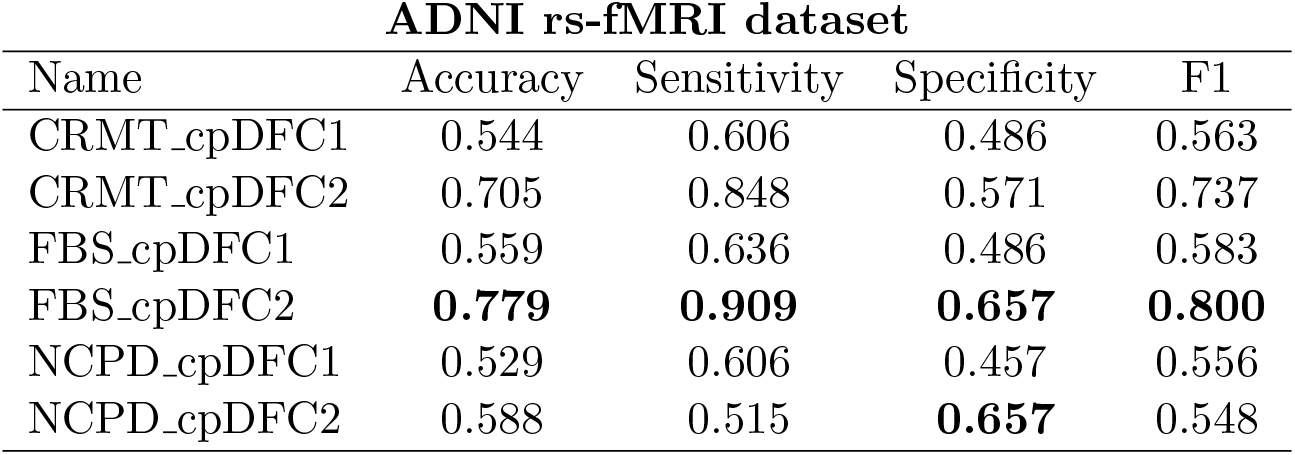
Classification results of CRMT, NCPD, and FaBiSearch change point methods on the ADNI rs-fMRI data.

| ADNI rs-fMRI dataset |  |  |  |  |
| --- | --- | --- | --- | --- |
| Name | Accuracy | Sensitivity | Specificity | F1 |
| CRMT_cpDFC1 | 0.544 | 0.606 | 0.486 | 0.563 |
| CRMT_cpDFC2 | 0.705 | 0.848 | 0.571 | 0.737 |
| FBS_cpDFC1 | 0.559 | 0.636 | 0.486 | 0.583 |
| FBS_cpDFC2 | **0.779** | **0.909** | **0.657** | **0.800** |
| NCPD_cpDFC1 | 0.529 | 0.606 | 0.457 | 0.556 |
| NCPD_cpDFC2 | 0.588 | 0.515 | **0.657** | 0.548 |

**Figure 6:**
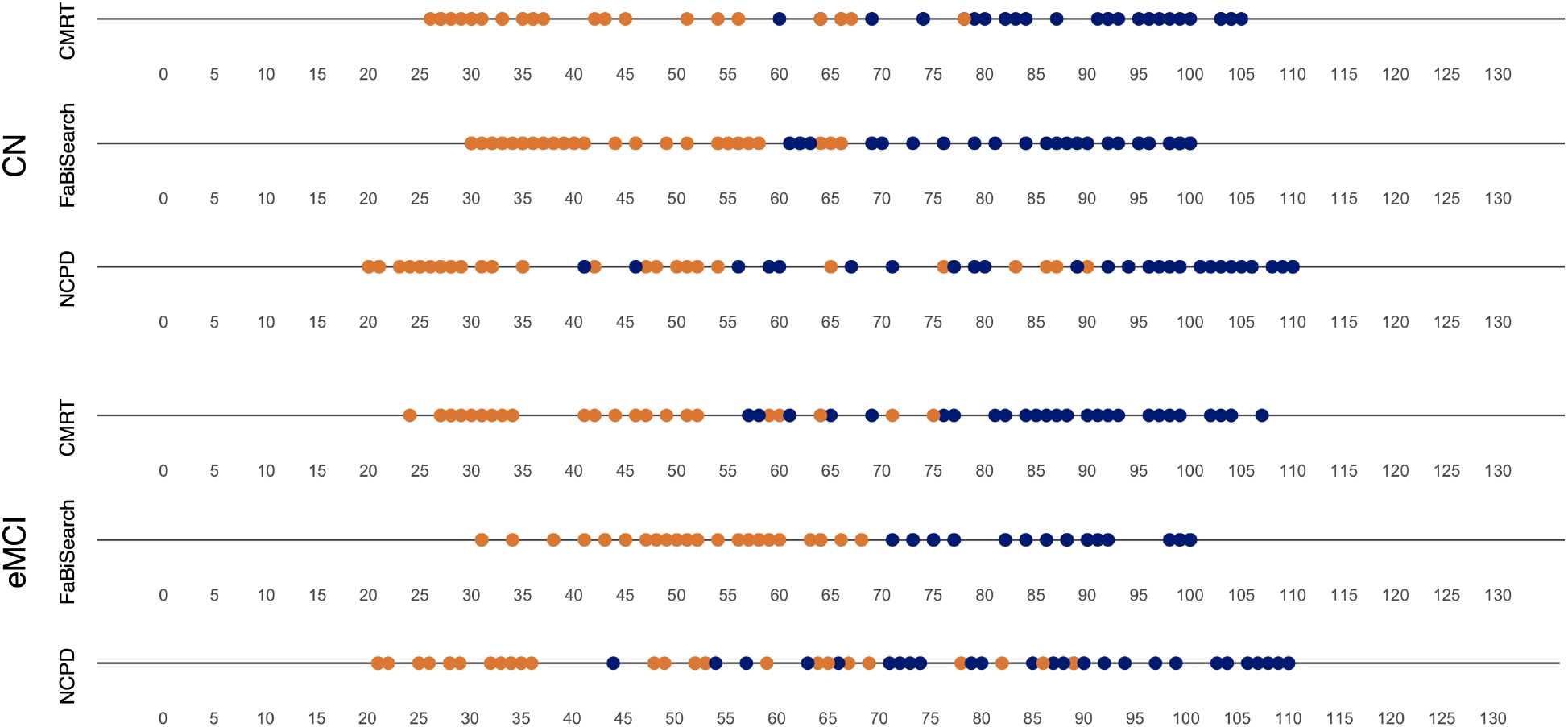
The detected change points for change point detection methodologies (CRMT, FaBiSearch, NCPD) for the ADNI rs-fMRI data. Orange and blue are used to denote the first and second detected change points, respectively.

### 5.4 Second MCI data set results

We present the classification results from the Mascali et al. (2015) dataset. Figure 7(top panel) shows the classification results using SFC, swDFC and cpDFC methods. Similar to the ADNI data set, FaBiSearch with two change points has a superior performance compared to the other methods. In addition, it appears that change point methods outperform swDFC methods. In Figure 7(bottom panels), we again find that there is no particular patterns between the different combinations of step size and window size and classification performance. The best performing models have a window size of approximately 40 and step size between 5 and 20 depending on the evaluation metric. For swDFC, performance on this dataset mirrors ADNI but is often inferior. Given the smaller size of this study (*n* = 20) compared to the ADNI data set (*n* = 68), it is possible that the performance discrepancies between the combinations of window and step size are further exacerbated in smaller sample size settings.

**Figure 7:**
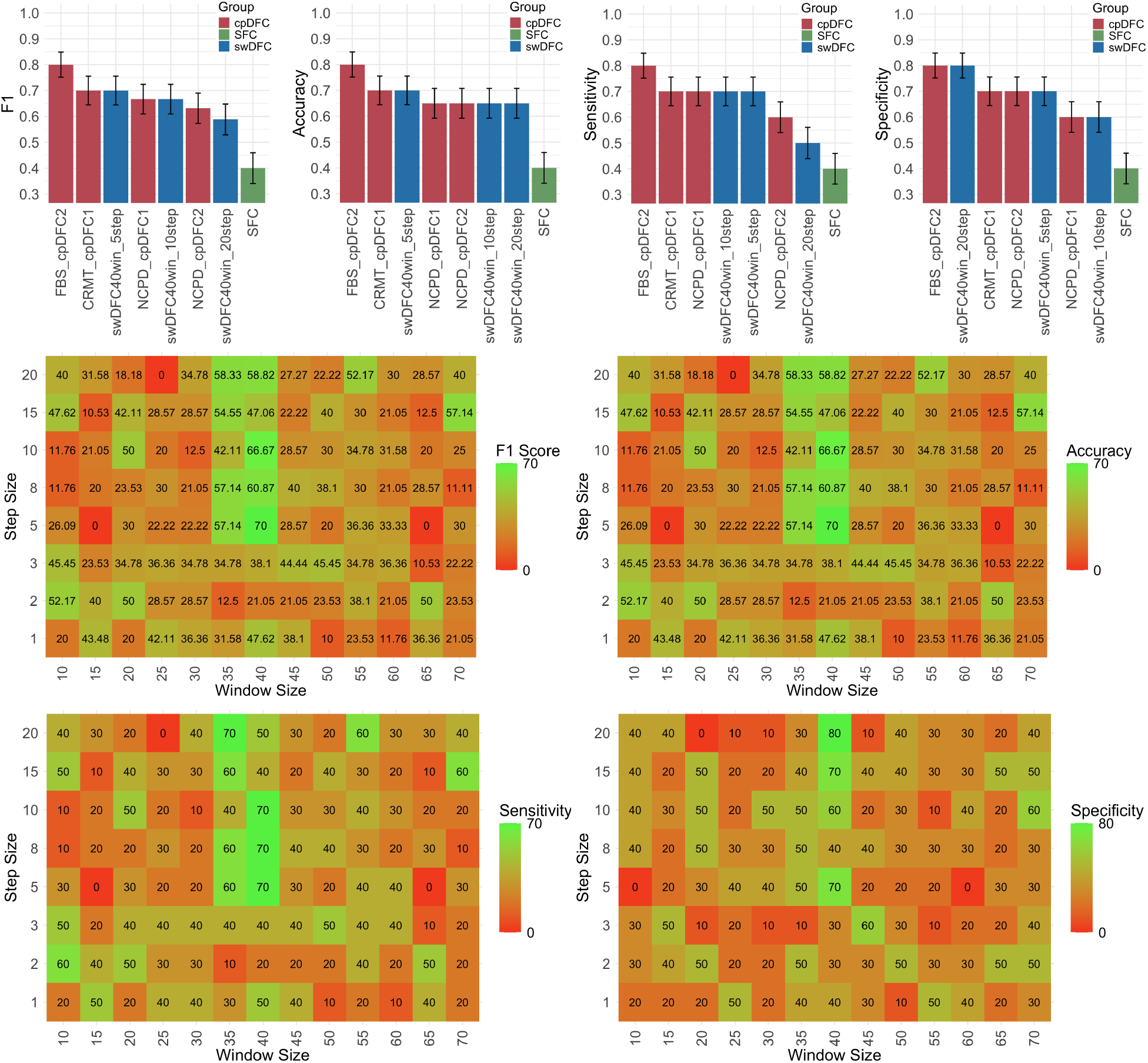
The classification results of CN subjects and subjects with MCI from the Mascali et al. (2015) rs-fMRI dataset. (Top panel) F1, accuracy, sensitivity, and specificity results of all methods. SFC, cpDFC, and swDFC correspond to static, change point, and window based dynamic functional connectivity, respectively. (Bottom panels) Heatmaps of F1 score, accuracy, sensitivity, and specificity. Results are shown across window sizes [10, 70] in increments of 5 and step sizes [1, 2, 3, 5, 8, 10, 15, 20].

We also consider other change point detection methodologies. Figure 8 shows the detected change points for both CN and subjects with MCI for the Mascali et al. (2015) data set for CRMT, FaBiSearch and NCPD. Table 2 shows the classification results for these change point detection methods. For all evaluation criteria, we find that FaBisearch with two change points, again, has the best performance. CRMT with one change point has the next best performance, and then both NCPD results. CRMT with two change points has the worst performance.

**Table 2:**
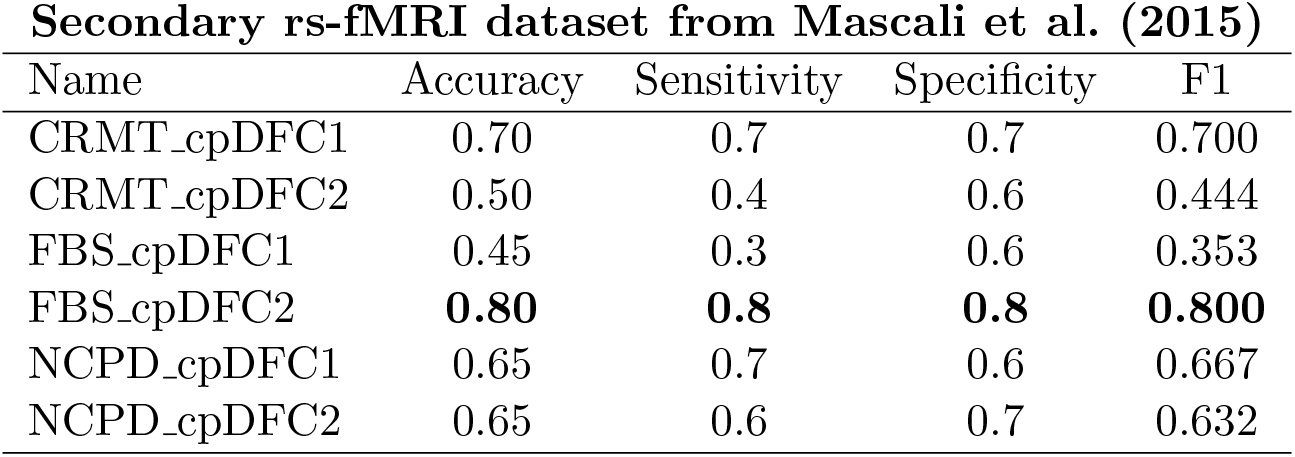
Classification results of CRMT, NCPD, and FaBiSearch change point methods on the Mascali et al. (2015) rs-fMRI MCI data.

| Secondary rs-fMRI dataset from Mascali et al. (2015) |  |  |  |  |
| --- | --- | --- | --- | --- |
| Name | Accuracy | Sensitivity | Specificity | F1 |
| CRMT_cpDFC1 | 0.70 | 0.7 | 0.7 | 0.700 |
| CRMT_cpDFC2 | 0.50 | 0.4 | 0.6 | 0.444 |
| FBS_cpDFC1 | 0.45 | 0.3 | 0.6 | 0.353 |
| FBS_cpDFC2 | **0.80** | **0.8** | **0.8** | **0.800** |
| NCPD_cpDFC1 | 0.65 | 0.7 | 0.6 | 0.667 |
| NCPD_cpDFC2 | 0.65 | 0.6 | 0.7 | 0.632 |

**Figure 8:**
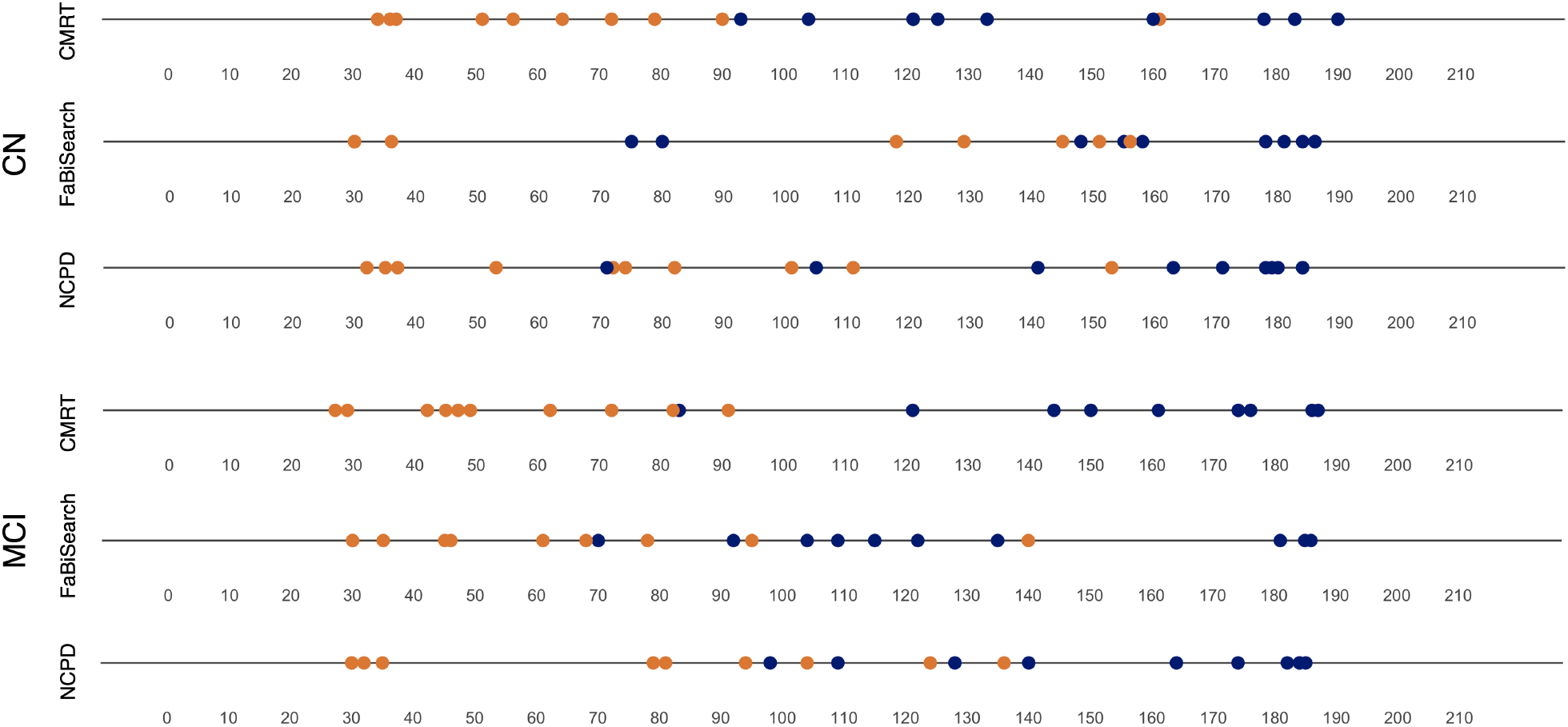
The detected change points for change point detection methodologies (CRMT, FaBiSearch, NCPD) for the Mascali et al. (2015) rs-fMRI MCI data. Orange and blue are used to denote the first and second detected change points, respectively.

### 5.5 Ensemble analysis

We present the results of the post-hoc ensemble analysis in Figure 9. As a reference, we also include the results from the FBS cpDFC2 model, which had the best stand-alone classifier performance. Figure 9(a) shows there is a clear upward trajectory as we combine more models. However, there is a diminishing return effect. The best accuracy (91.17%) was obtained by the ensemble method that combined the best eight models based on their highest F1 score (top-8 model). This model had the best performance across all accuracy metrics, except sensitivity, where the best model combined the best ten models based on the highest F1 score (top-10). However, the top-8 model had the best performance (91.42%) in terms of the F1 score, which balances both sensitivity and precision. This indicates that even though the top-10 ensemble was superior in terms of sensitivity, this was at a cost to precision. The top-8 ensemble model had a superior performance compared to all single condition models, including the best, FBS cpDFC2 (*p* = 0.006).

**Figure 9:**
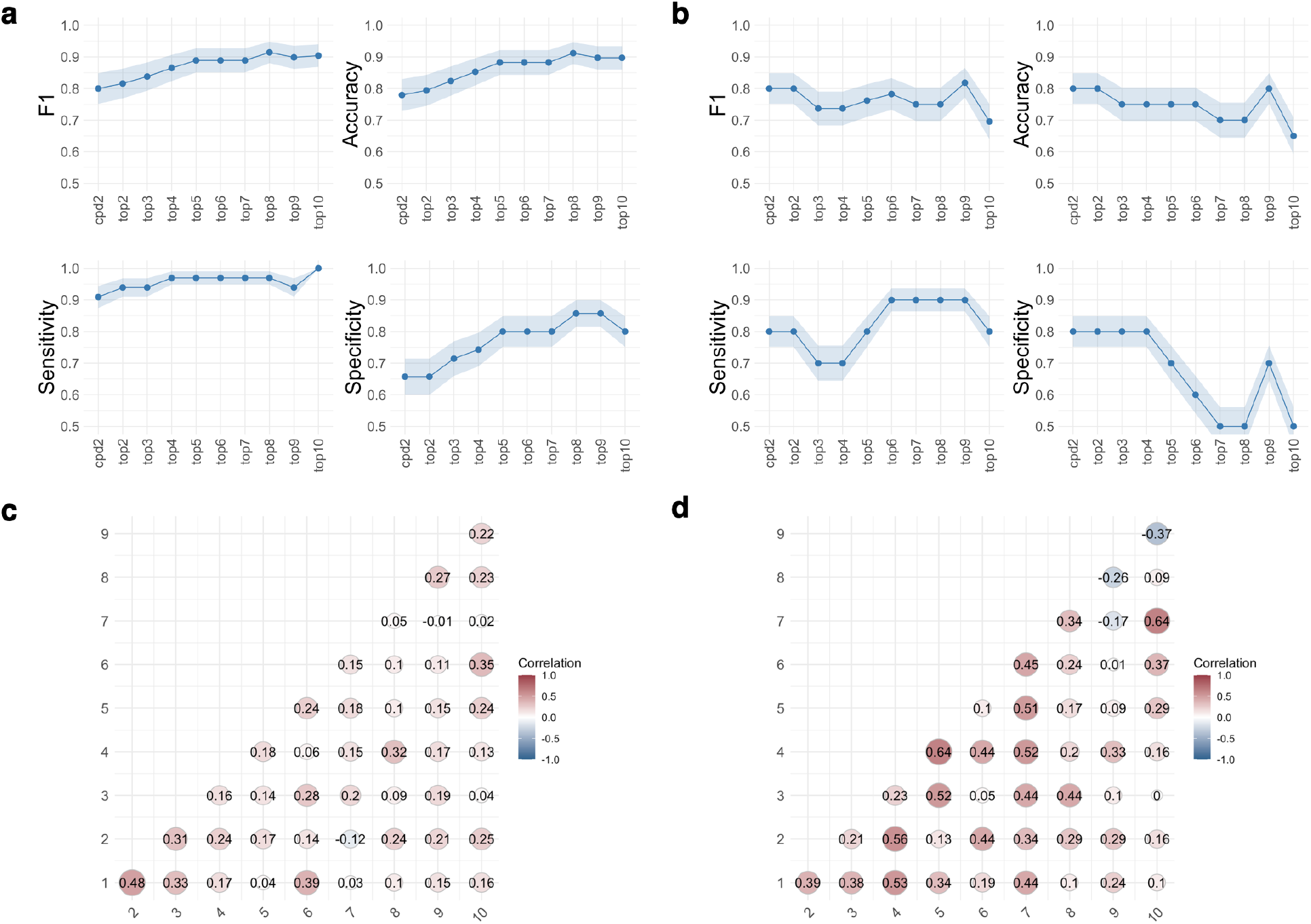
The ensemble model results from the classification study of CN subjects and subjects with eMCI. Panels (a) and (b) show the ensemble results for the ADNI and Mascali et al. (2015) rs-fMRI datasets, respectively. For the ensemble model, we combined the predictions from the top-2 to the top-10 classifier models, as determined by their highest F1 score. The FBS cpDFC2 calculated as 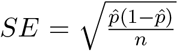. (c) and (d) the correlation between predicted probabilities of eMCI model is the best stand-alone classifier. Shaded regions indicate ± the standard error of proportion across the top-10 classifiers, as determined by their highest F1 score for the ADNI and Mascali et al. (2015) data sets, respectively.

Figure 9(c) shows a correlation plot of the predicted probabilities for the top-10 models determined by the highest F1 score. The predicted probabilities of the best model (cpDFC2) and the second best model (swDFC40w 5s), and the best model (cpDFC2) and the sixth best model (swDFC35w 20s) were significantly correlated (*t*-test, *α <* 0.05, adjusted for multiple comparisons using Benjamini and Hochberg, 1995). Hence, we can conclude that some of the models in the ensemble method are related and may be adding some of the same prediction power in the classification task. Figure 9(b) shows the same ensemble analysis for the Mascali et al. (2015) dataset. The lack of a monotonically increasing performance pattern as more models are combined may stem from the small sample size. Alternatively, it could be due to the larger number of higher correlations between the models (Figure 9(d).

## 6 Discussion

We begin our discussion by considering the cpDFC results, which show that there is group-level similarity in the locations of the change points between the CN subjects and subjects with eMCI (Figure 2). Since subjects are not given explicit directions during the experiment, they are allowed to flow between states at any rate. With the lack of external stimulus, each subject’s patterns of thoughts, and consequently functional brain dynamics, are unconstrained. Our results indicate that the rate of change between states is unique for each subject and does not seem to provide any information for differentiating CN and subjects with eMCI. Furthermore, the uniqueness of the rate of change between time segments emphasizes the high individual variability in temporal brain dynamics. This variability could be due to differences in thought patterns, mental states, and underlying neural mechanisms that are not externally directed due to the resting-state nature of the experiments (Gonzalez-Castillo et al., 2021, 2019; Vidaurre et al., 2017). cpDFC methods applied at the subject level then provides a way to account for these individual discrepancies in a systematic and statistically sound manner. Although no group-wise differences were identified in terms of change point locations, there appears to be a group-wise difference for the first change point detected. We found that the first change point for the CN group was on average earlier than for the eMCI group. This may suggest that CN subjects settle more quickly into stable homeostatic patterns than their eMCI counterparts. In contrast, part of the cognitive impairment seen in eMCI could be related to the slower transition to resting states.

Next, we consider the single model results, which suggest that, in general, DFC methods provide information that is useful for eMCI classification. However, deciding the exact combination of parameters in the swDFC method that have the best classification results is not a clear task. Hence, multiple change point detection such as FBS cpDFC2 provides a clear advantage. One distinction between swDFC and FBS cpDFC2 is that the former uses overlapping segments, which may be able to better capture “smooth” transitions between states, although this does not seem to significantly impact the classification of eMCI. It leads to many windows, which have overlapping and therefore redundant information that may not be useful for differentiating subjects. Furthermore, it also leads to a higher-dimensional classification problem, since the number of time segments is larger, hence the feature vector for each subject is inherently larger. In a sample-constrained setting, such as fMRI, this causes issues in terms of dimension reduction and variable selection. Another important distinction between the two methods is that the rate of change is estimated independently for each subject in FBS cpDFC2, whereas in swDFC it is assumed to be the same rate (the same window and step size is used across all subjects). This suggests that even though the rates of change between subjects may be different, the transition between states occurs in an ordinal manner. Furthermore, the FBS cpDFC2 method outperformed the swDFC method, suggesting that finding subject-level differences in the rate of temporal change is an important consideration for differentiating groups of subjects. These observations suggest that each subject has a unique “temporal fingerprint” that interacts closely with their state, as well as classification between CN and subjects with eMCI.

For swDFC, we have two conclusions. First, finding the correct choice of swDFC parameters is a non-trivial task. This is evident in the heatmaps (Figures 4, 7), which do not show a clear pattern or relationship between the choice of window and step size. Furthermore, the results for swDFC are not always superior to those of SFC. There are several instances of window and step size combinations where swDFC leads to inferior results to the null model. This appears to be related to the instability of the swDFC measures, partly due to the characteristics intrinsic to sliding windows, but also due to individual differences in brain dynamics. Second, the results of FBS cpDFC2 may not be statistically significant different to the best combinations of swDFC, but it has a superior performance to the average of all combinations of swDFC: the accuracy of the swDFC ranges from 0.22 to 0.72, and the group average in all swDFC methods (0.56) was not statistically significant different to SFC (0.57). This suggests that without knowing the precise window and step sizes to use, which is not possible *a priori* in a setting such as fMRI, the use of swDFC may lead to poor and unstable results. Furthermore, it requires testing a large number of combinations of window size and step size, and therefore hypotheses, which can inflate Type I error and false discovery rate. It also reinforces the need to select the appropriate window and step sizes for accurate and robust results. In fact, this may explain some of the issues that seem to be related to instability in DFC studies and their results (Lurie et al., 2020).

The performance of FBS cpDFC2 is at least as good as the best swDFC also supports the notion that change point detection is similar to finding the optimal window size, which has the downstream effect of providing estimates of dynamic networks with greater stability. This suggests that change points can be used to directly model the dynamical processes in brain networks, rather than requiring a grid search over all possible combinations, and that this is generally more informative for differentiating subjects. In this way, we can use change point detection as a model to find meso-scale patterns for a time-dependent, stochastic process.

We demonstrate the stability of the cpDFC model by considering the number of unique features selected in the folds. FBS cpDFC2 ranks close to the best in terms of selecting the smallest number of unique features, which supports the notion that cpDFC is useful for recovering both stable features and features that are relevant to differentiate CN and subjects with eMCI.

Our results from the outputs of the FBS cpDFC2 model show that the most predictive ROIs for eMCI occur within the frontal, temporal, cerebellum, occipital, and parietal communities. Furthermore, these features are generally from windows in the latter part of the experiment, in particular the 3^*rd*^ (final) time window. This suggests that the most informative features of brain dynamics, when differentiating between CN subjects and eMCI subjects, arise from the later stages of an rs-fMRI experiment and may not be related to the initial states as the subjects begin the experiment in the scanner. For example, different subjects may initially have different thought patterns, levels of alertness, or mental drift states, which may be reflected in differing functional organization. However, it seems that later in the experiments, subjects converge to a more common steady state, which has important information related to differentiating between CN and subjects with eMCI. Furthermore, we found our most informative features from local characteristics in a few brain regions, which suggests that strong dynamic predictors are concentrated in key regions, rather than being dispersed across the entire brain.

There are also interesting patterns in the features and localization in the brain (Figure 5). In the frontal lobe, the same two features, clustering coefficient and local efficiency, were chosen for two nodes, 6 and 73. Since the frontal lobe is key in executive functioning and is implicated in many brain disorders (Miller, 2007), its activity may serve as an important indicator of eMCI and possibly the onset of AD (Risacher et al., 2009; Tijms et al., 2013). The difference in mean values between the CN and eMCI groups for these nodes is minimal, which may suggest the following. First, small changes in the local efficiency of information transfer and clustering density may be sufficient to differentiate between CN and eMCI. In relation to this, there are likely subtle differences in the function of the frontal lobe, as is evident through executive function, which is important and relevant given the lack of clear behavioral differences between the CN and eMCI groups. Second, the general notion that the efficiency and clustering in AD are reduced in these regions may not hold for eMCI, as the clustering coefficient in nodes 6 and 73, and the local efficiency of node 6 actually increased in the eMCI group (Tijms et al., 2013).

The most informative feature was the degree of node 70’s in time segment 2, and it showed a decrease (on average) in CN subjects compared to the subjects with eMCI. From a graph-theoretic perspective, this node saw a decrease in functional connections in subjects with eMCI. As a member of the parietal lobe, node 70 is crucial and involved in many cognitive functions such as attention, planning, and information processing (Culham and Kanwisher, 2001; Husain and Nachev, 2007; Silver and Kastner, 2009). Furthermore, the parietal lobe has also been studied in the context of AD, where its atrophy and reduced metabolic capacity have been associated with the disease (Buckner et al., 2008; Minoshima et al., 1997; Karas et al., 2004).

The shortest path feature of nodes 107 and 111 in the cerebellum was also an important feature. For both nodes, there was a large increase (on average) in the length of the shortest paths that connect these nodes for the subjects with eMCI compared to the CN subjects. The cerebellum, which is best known for its role in motor coordination (It ō, 1984) and is not typically involved in AD, has recently been studied in the context of the onset of the disease (Jacobs et al., 2018). In addition to motor control, the cerebellum acts as an important hub and transmits to other brain regions; therefore, its relevance to eMCI may be linked to its capacity as a functional transmitter for cognitive and information processing abilities (Ramnani, 2006; Strick et al., 2009; Schmahmann, 2019). To compute the shortest paths, we first need to estimate the correlation and then apply a cutoff, therefore, we can also have the following interpretation; the number of strong (|*r*| *>* 0.5) pathways between nodes through the cerebellum decreased markedly in subjects with eMCI.

The differences in the top selected features in the classification study of CN subjects and subjects with eMCI are non-obvious. For example, betweenness centrality for the cerebellum ROI is higher in the eMCI group, but lower in the ROIs of the occipital and temporal lobes. Another example is the local efficiency for nodes 6 and 73; in the former, the eMCI group has larger values, but for the latter, the opposite is true.

We also included an ensemble study. Our simple strategy of taking the average prediction probabilities from several of the best models to generate an ensemble model was sufficient to improve the prediction performance. This improvement over any single model is also comparable to other more highly parameterized and non-linear models from other studies (see Du et al., 2018 for comparisons). In particular, the top-8 model was significantly better than the best single model, FBS cpDFC2. Although prediction accuracy is desirable, there is an alternative narrative; combining the swDFC and cpDFC methods, each of which captures different time scales, is beneficial for the eMCI and CN classification, and important information is embedded as a multiscale process. The results of the ensemble model are also supported by analyzing the correlation between the predicted probabilities of each classifier. We discovered that, in general, the predictions in the top-10 models were not highly correlated except between the FBS cpDFC2 and swDFC40w 5s models, as well as between the FBS cpDFC2 and swDFC35w 20s models. This finding further supports the idea that models with different timescales provide discrete information that is complementary to subject classification. This conclusion is intuitive, as we typically assume that the brain is plastic, dynamic, and evolving. On one end of the time scale there may be fast neural impulses that change in an instant. On the other end of the time scale there may be the slow, evolutionary process of learning. From the brain activity that is captured using fMRI, there must be multiple different types of dynamics that occur at different time scales, and our findings support this notion. However, it is not clear how to determine which time scales are important and how best to incorporate this information. It may also be useful to think of change point detection models as important for uncovering mesoscale dynamic structure in rs-fMRI experiments; SFC provides a stationary picture, while windows provide a microscale, overlapping dynamic structure. These ideas have been presented before in the work of Betzel and Bassett (2017), however, our study is the first to explore this phenomenon in eMCI detection in an rs-fMRI study.

The improvement in performance in the ensemble models shows evidence of diminishing marginal returns. In other words, the performance of adding more models to the ensemble models improves (the best performance occurs at the top-8 ensemble); however, afterward the performance stabilizes and begins to drop off thereafter. This suggests that improvements in classification performance through ensemble models require each prediction from different timescales to provide unique information that is not present in the other models with different timescales.

Many studies have explored the classification of subjects with eMCI and CN subjects using ADNI rs-fMRI data (https://adni.loni.usc.edu/). Typical classification accuracy results range from approximately 70% to 90% depending on the modality (genetic, behavioral, structural, functional) and the type of classifier model (Suk et al., 2016; Wee et al., 2013). A key observation from these studies is that combining modalities and information at different levels within a modality has the greatest enhancing effect on performance. Using a simple ensemble method of different time scales, we achieve state-of-the-art performance with fewer features and with only a single modality compared to other methods which often utilize other modalities in addition to rs-fMRI. Studies of other types of disease and methodological approaches support these observed trends (Du et al., 2018). Similarly, we find an improvement in classification metrics by taking a multi-timescale approach in our post-hoc ensemble analysis. Clearly, predicting classes of neurodegeneration from measurement modalities is non-trivial, hence it makes sense to integrate features from different modalities as well as scales within modalities to get a more diverse and cohesive set of predictive features.

Lastly, we provide additional results from a secondary MCI classification study (Mascali et al., 2015) and apply different change point detection techniques (Cribben and Yu, 2017; Ryan and Killick, 2023), to further support our main results. These secondary results show that, similar to the main results, FaBiSearch has a superior performance to swDFC and other change point methods. We further show that the improvement from the ensemble models holds in this secondary study, albeit with fewer ensemble models required to obtain optimal results. Given the differences in data collection and pre-processing methods, these secondary results suggest that our main findings are consistent and robust across other data generation schemes. There are also vast differences in prediction performance based on the type of change point detection method used. However, unlike swDFC, we did not observe such a wild discrepancy between the best and the worst performers. It appears that all together, cpDFC is at least as good as SFC unlike swDFC which oftentimes leads to inferior results. Furthermore, the classification performance suggests that the combination of change points in the low-rank graph structure detected by FaBiSearch as well as the modeling of time series through NMF may be important for downstream tasks. In addition, the multiple change point method, CRMT, appears to perform slightly better than NCPD, and closer to FaBiSearch. Given CRMT initially projects the time series data into a lower dimensional space using truncated SVD, the way we set up change point detection for this method implies that it detects changes in low-rank structure akin to FaBiSearch. Given previous work which looks at the low-rank projection of neural activity based on stimulus and individual characteristics (Yatsenko et al., 2015), this may explain the improved efficacy over NCPD.

### 6.1 Limitations

An important limitation of our study is the small sample size of the rs-fMRI experiments. Given the cost and labor associated with collecting rs-fMRI data, it is quite typical for these studies to have small samples. Ultimately, this reduces the generalizability of the results as the statistical power is limited. With this in mind, it is difficult to conclude that predictive models will have a similar performance with a large sample size. However, our results suggest that even with this limitation, change point detection and FaBiSearch in particular is an important statistical tool for finding stable, consistent patterns from high-dimensional multivariate time series data. Furthermore, the statistical properties and patterns captured by FaBiSearch provide valuable information on the mechanisms of disease that are important for the detection of eMCI.

## 7 Conclusion

In this work, we compare the performance of SFC, swDFC, and cpDFC for an eMCI classification task using a single modality, rs-fMRI data. First, we show that multiple cpDFC is superior to swDFC based methods for disease classification. Second, we show that DFC is more useful than SFC for the classification of a difficult task such as eMCI detection. Third, we delineate several key advantages of cpDFC in comparison to swDFC that make it an attractive approach to capture DFC beyond the downstream classification task. Finally, using a simple ensemble method of different time scales, we achieve state-of-the-art performance with fewer features and with only a single modality compared to other methods which often utilize other modalities besides rs-fMRI. Together, our results demonstrate the power of change point detection with FaBiSearch for a difficult classification task between subjects with eMCI and CN subjects. Our work suggests that change point detection is important in the statistical toolbox for analyzing fMRI data, and is valuable in uncovering hidden disease dynamics in rs-fMRI experiments. We hope this study inspires a renewed interest in advancing research on change point detection techniques aimed at estimating dynamic functional connectivity.

Although in this study we focus on FaBiSearch based change point detection for eMCI classification, there is a rich literature on change point detection in multivariate time series, a wide array of other neurological disorders, and even variability in cognitive processing. Future works could look at the potential differences in change point detection approaches depending on the context and downstream prediction task. Additionally, it remains unclear how best to select the time scales that are the most important for downstream tasks and how this interacts with individual differences and outcomes. Although our approach of concatenating feature vectors together worked well in this context, future studies could explore how to select which time scales are most important and how to best combine the different levels of information they provide.

## 8 Data and code availability statement

Due to the sensitive nature of the data used in this study, as well as the terms of use for both sources, we are unable to directly share the data used. The ADNI and Mascali et al. (2015) datasets used were derived from the following public domains http://adni.loni.usc.edu/ and https://dataverse.harvard.edu/dataverse/restAD, respectively. All `R` code implementing experiments is available on GitHub.

## 9 Acknowledgements

This research was enabled by support from WestGrid (www.westgrid.ca) and the Digital Research Alliance of Canada (www.alliancecan.ca). The first author was supported by the Alexander Graham Bell Canada Graduate Scholarship (CGS M) from the Natural Sciences and Engineering Research Council of Canada (NSERC), the Alberta Innovates Graduate Student Scholarship (AIGSS) from Alberta Innovates, the Alberta Graduate Excellence Scholarship (AGES) from Alberta Advanced Education, the Richard B. Stein Neuroscience Graduate Studentship from the Neuroscience and Mental Health Institute of Alberta (NMHI), and the H. Jean McDiarmaid Scholarship from the University of Alberta Faculty of Medicine and Dentistry. The second author was supported by the NSERC Discovery Grant RGPIN-2024-06102.

## 10 CRediT author contribution statement

**Martin Ondrus:** Conceptualization, Methodology, Software, Validation, Formal analysis, Resources, Writing – original draft, Writing – review & editing, Visualization.

**Ivor Cribben:** Conceptualization, Methodology, Software, Validation, Formal analysis, Resources, Writing – original draft, Writing – review & editing, Visualization.

## 11 Ethics

This article complies with the ethical guidelines of the MIT Press, COPE, and ICMJE. The research presented has been conducted and reported with integrity, transparency, and accountability. No specific ethical issues or conflicts of interest are present in this submission.

## A Appendix

### A.1 Algorithms for time-varying functional connectivity estimation

#### Algorithm 1

Algorithm for estimating dynamic functional connectivity using sliding windows.

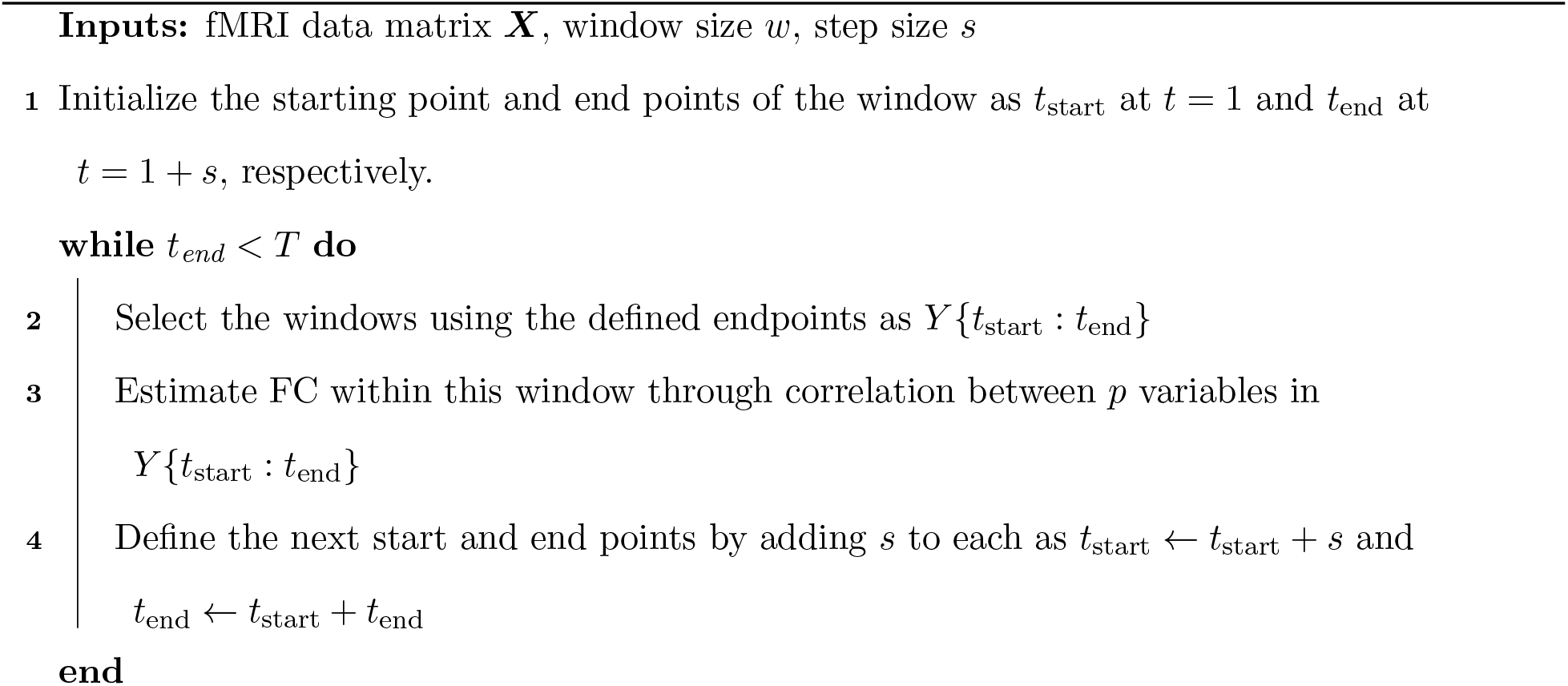

#### Algorithm 2

Change point detection algorithm using FaBiSearch which can be applied recursively to find multiple change points.

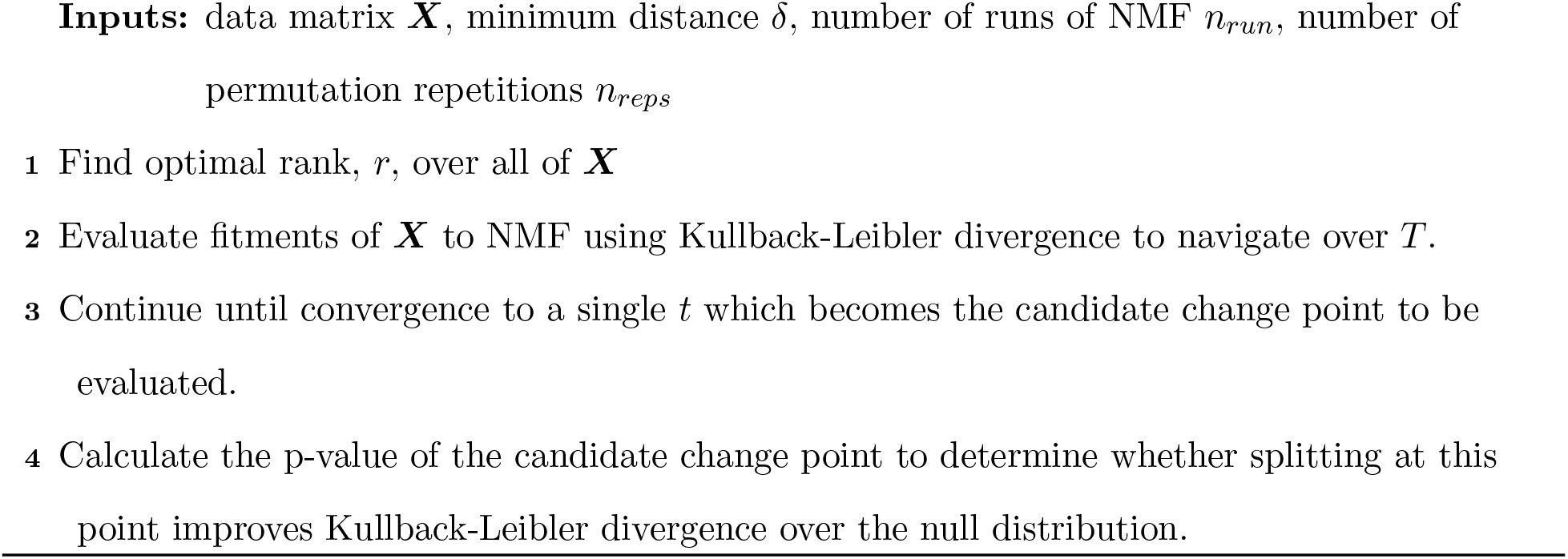

### A.2 Graph Theoretic Features

#### Degree

The degree of a node can be calculated from the following definition:

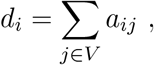

where *d*_*i*_ ∈ ℕ_0_ is the degree of the node *i, V* is the set of all nodes, and *a*_*ij*_ is the intersection of the nodes *i* and *j* in the adjacency matrix.

#### Clustering coefficient

From Wasserman and Faust (1994), it is given by:

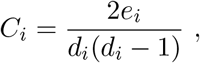

where *C*_*i*_ ∈ [0, 1] is the clustering coefficient for node *i, d*_*i*_ is the degree of node *i*, and *e*_*i*_ is the number of edges between node *i* and neighbors *d*_*i*_.

#### Shortest path

We use the notation ℓ_*ij*_ ∈ ℕ_0_ as the fewest number of edges that connect nodes *i* and *j* together to define the shortest path. The shortest path can be calculated using different algorithms, although we use breadth-first search (BFS: Cormen et al., 2022) in our implementation.

#### Degree assortativity

Newman (2002) define degree assortativity as

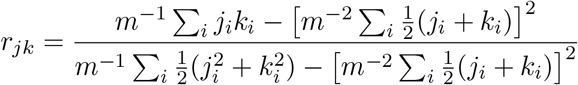

where r ∈ [−1, 1] is the degree assortativity of the graph, m is the total number of edges, and j_i_ and *k*_*i*_ are the degree of the nodes *j* and *k* that are connected through *i*.

#### Local efficiency

The definition of local efficiency from Latora and Marchiori (2001) is

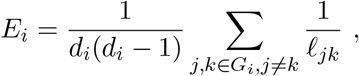

where *E*_*i*_ ∈ [0, 1] is the local efficiency measure of node *i, d*_*i*_ is the degree of node *i*, and ℓ_*jk*_ is the shortest distance between nodes *j* and *k* in the sub-graph of 𝒢_*i*_.

#### Betweenness centrality

The definition of betweenness centrality (Freeman, 1977) follows:

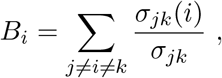

where *B*_*i*_ ∈ [0, 1] is the betweenness centrality of node *i, σ*_*jk*_ is the total number of shortest paths between nodes *j* and *k*, and *σ*_*jk*_(*i*) is the total number of shortest paths between *j* and *k* that pass through *i*.

### A.3 Evaluation metrics

We evaluated the performance of our models from the classification study using the following four metrics:

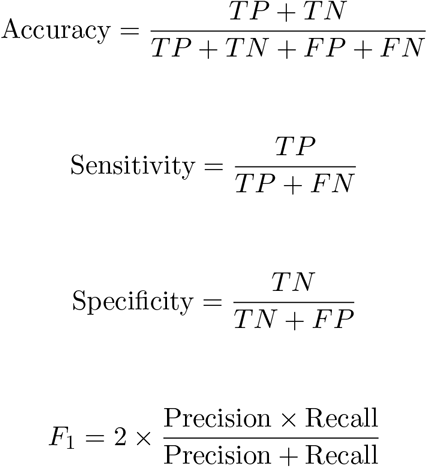

where Precision 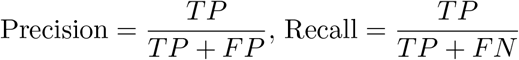 and,

*TP* denotes True Positives; The number of correctly labeled eMCI.

*TN* denotes True Negatives; The number of correctly labeled control.

*FP* denotes False Positives; The number of control incorrectly labeled as eMCI.

*FN* denotes False Negatives; The number of eMCI incorrectly labeled control.

### A.4 Additional results

As the results in the main article show, there is a concentration of strongly differentiating features in the later time segments (Figure 5) for the classification task. This is highlighted also in Figure A1, especially in the ROIs close to the main diagonal, and also in the squared difference row, which are the largest for time segments 2 and 3. It is evident that the group-wise characteristics become more stable and contribute most to the differentiation in later time windows. Furthermore, it is clear from Figure A1 that global changes in graph structure are subtle, since the averaged adjacency matrices are generally relatively similar, but localized differences in functional connectivity have widespread effects related to the global graph structure. For example, we observed differences in the shortest path metric, which can be strongly influenced by changes in one or a few edges, especially if there is a lack of redundancy in connections between nodes. Therefore, these subtle differences in the graph structure have strong consequences in distinguishing CN from patients with eMCI.

As we restrict each subject to two change points, each subject has 3 stationary segments, or partitions between change points (or stationary modes). We can then compare group-wise differences in the adjacency matrices between consecutive segments. We calculate the average adjacency matrix for each segment and class (CN or eMCI), using the following procedure. For each group (CN: *n* = 35, eMCI: *n* = 33), 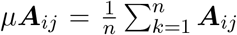. We can then calculate the entry-wise difference between the groups using *µ****A***^diff^ = *µ****A***^CN^ − *µ****A***^eMCI^. Figure A1 (rows 1 and 2) shows the average adjacency matrices grouped by segment (1, 2, or 3) and class (CN or eMCI), while Figure A1 (row 3) shows the differences in class between segments for all 120 ROIs. The differences appear to concentrate close to the diagonal of the adjacency matrices. This difference is especially pronounced between ROIs 50-60. In addition, subjects with eMCI have stronger off-diagonal connections, notably at the intersection of ROIs 10-20 and 80-90.

**Figure A1:**
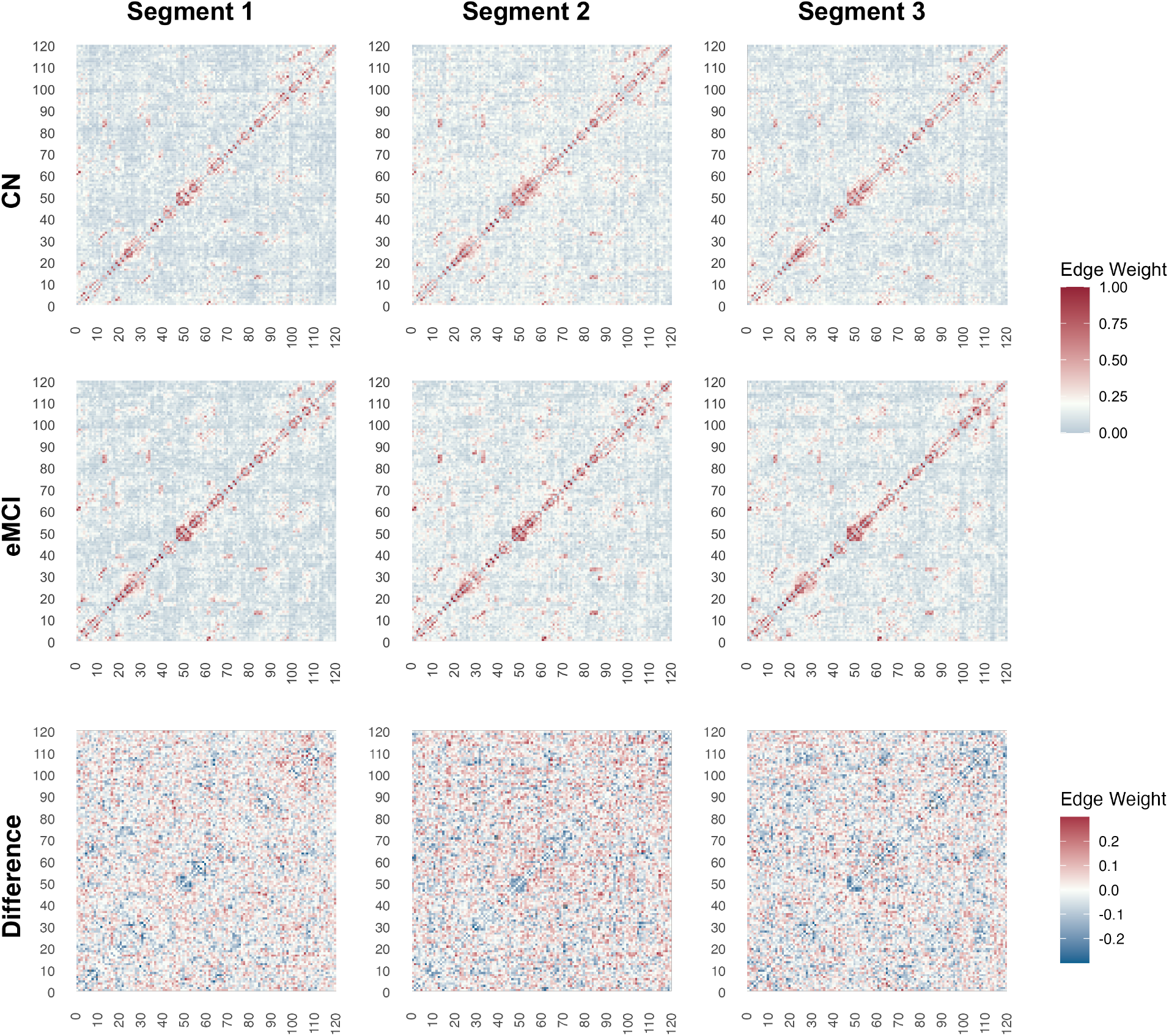
Averaged adjacency matrices across temporal segments (columns), and subject class (rows) for the FaiSearch (cpDFC2) method in the classification study of CN subjects and subjects with eMCI from the ADNI rs-fMRI dataset. Node labels (1 to 120) correspond to the AAL atlas (Tzourio-Mazoyer et al., 2002) labels. The difference between the CN and the eMCI adjacency matrices computed for each segment is shown in the third row.

**Figure A2:**
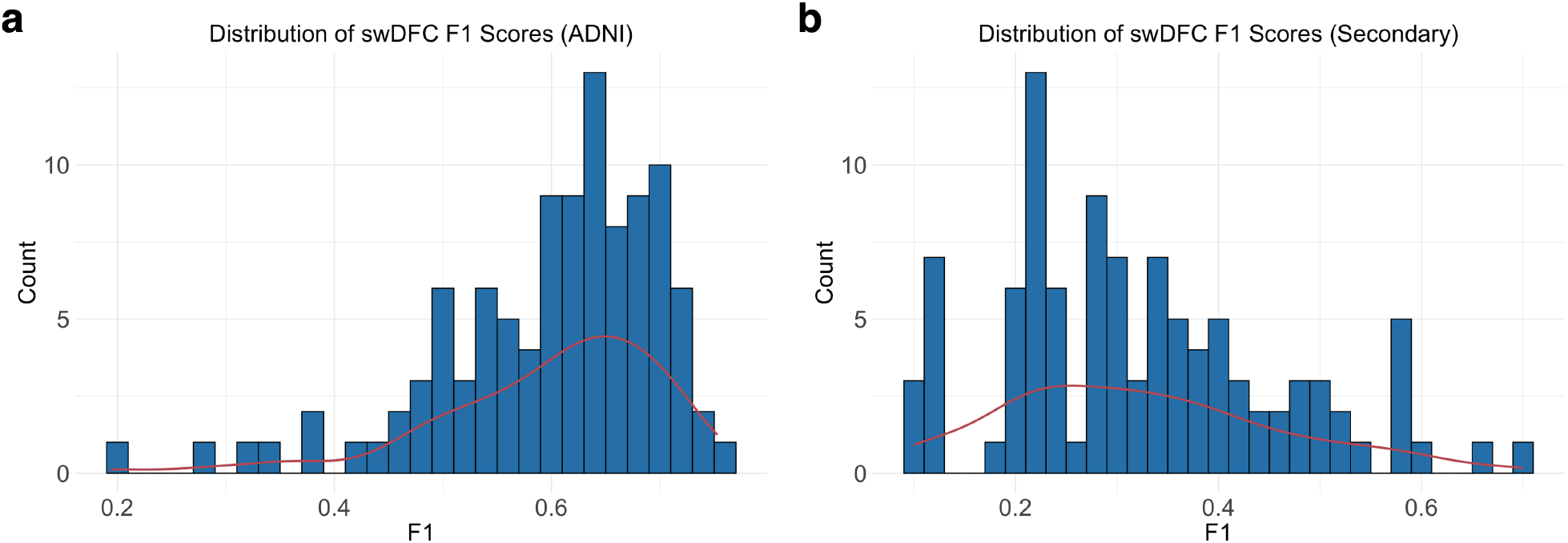
Histograms of the swDFC F1 scores for the (a) ADNI, and (b) Mascali et al. (2015) rs-fMRI datasets, where the red lines corresponds to the kernel density estimate.

**Figure A3:**
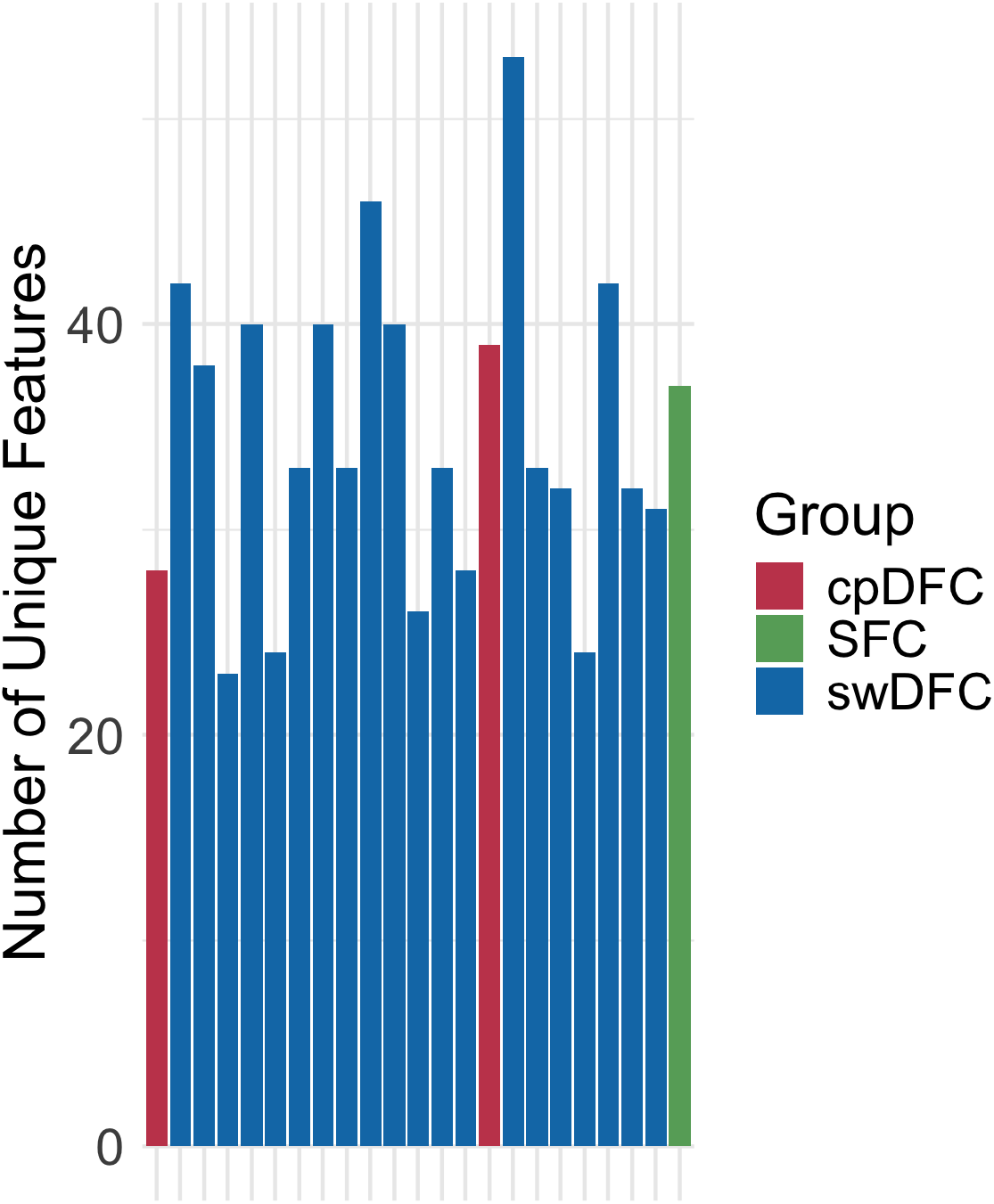
A bar plot of unique features selected across folds for cpDFC, swDFC, and SFC. Lower values indicate more consistent across fold selection of features. Bars are ordered left to right according to descending LOOCV F1 score for each method.

